# Spatiotemporal factors shape mosquito virus diversity and virome composition in a host and virus-dependent manner

**DOI:** 10.1101/2025.09.05.674492

**Authors:** Cassandra Koh, Lionel Frangeul, Hervé Blanc, Aurélien Gibaud, Annabelle Henrion-Lacritick, Rohit A Chitale, Sébastien Boyer, Jean-Bernard Duchemin, Philippe Dussart, Romain Girod, Nina Grau, Carine Ngoagouni, Anamarija Butkovic, Maria-Carla Saleh

**Affiliations:** Institut Pasteur, Université Paris Cité, Viruses and RNA Interference Unit, F-75015 Paris, France; Mayo Clinic, Division of Infectious Diseases, Jacksonville, FL 32224, USA; Institut Pasteur du Cambodge, Medical and Veterinary Entomology Unit, Phnom Penh 12201, Cambodia; Institut Pasteur, CNRS UMR2000, Ecology and Emergence of Arthropod–Borne Pathogens Unit, F-75015 Paris, France; Institut Pasteur de la Guyane, Vectopôle Amazonien Emile Abonnenc, Cayenne 97306, French Guiana; Institut Pasteur du Cambodge, Virology Unit, Phnom Penh 12201, Cambodia; Institut Pasteur de Madagascar, Medical Entomology Unit, Antananarivo 101, Madagascar; Institut Pasteur de Bangui, Medical Entomology Laboratory, Bangui BP 923, Central African Republic

## Abstract

Beyond the arboviruses they transmit to humans and animals, mosquitoes harbour diverse viral communities that shape infection and immunity dynamics. Understanding how these communities shift within mosquito populations across time and space has important implications for how we anticipate and manage arboviral disease epidemics. Here, we examined the viromes of rural and sylvatic mosquito species sampled longitudinally across two years from countries representing South America, Africa, and Southeast Asian geographies to quantify the relative contributions of host and ecological factors in shaping their virome dynamics. We found that virus diversity and virome composition are strongly influenced by host phylogeny and spatial factors and show limited temporal variation. By examining virus similarity networks across mosquito species and biogeographies, we observed frequent instances of within- and cross-genus *Culex*-*Anopheles* virus sharing in Madagascar. Taken together, these findings inform the development and implementation strategies of any virome-based vector control technology and raise new evolutionary questions on how host generalism shapes the spatial prevalence of viruses and their potential for disease emergence.

## INTRODUCTION

Mosquitoes harbour remarkably diverse virus communities, counting at least 48 families among them^1^, and metagenomics studies continue to uncover lineages of novel mosquito-associated virus^2^. Arboviruses, such as Zika, chikungunya, dengue or West Nile viruses, are only a small fraction of mosquito virosphere^3,4^. Phylogeny suggests that a large proportion of mosquito viruses are mosquito-specific, that is, have host ranges that are restricted to invertebrates. Their abundance and prevalence across natural mosquito populations makes them an integral part of the disease ecology of vector mosquitoes^4^.

Phylogenetic reconstructions also revealed that several arboviral lineages originated from mosquito-specific ancestors^5,6^. This has led to the view of arthropods, including mosquitoes, as sources of novel viral pathogens capable of jumping to vertebrate hosts^1,7^. Interestingly, some mosquito-specific viruses have been demonstrated to inhibit or enhance arboviruses in co-infected mosquitoes^8–14^. They can therefore influence mosquito vector competence through virus-virus and virus-host interactions, with multifaceted implications for mosquito-borne disease transmission.

Mosquito virus diversity and virome composition have been linked to host phylogeny^15–17^, yet the drivers of virome dynamics, that is, how virus diversity and virome composition vary across spatial and temporal scales, remain poorly explored. Understanding how host and spatiotemporal factors interact to shape mosquito viromes would support the estimation of arboviral disease emergence risk through surveillance and the development of virome-based vector control approaches.

Investigating spatiotemporal virome dynamics in natural mosquito populations requires studies that span large spatial and temporal scales, which are often logistically challenging^14,18,19^. While publicly available metagenomic datasets enable virome meta-analyses across broad geographies^4,20,21^, longitudinal datasets from the same sampling locations are rare. In addition, variation in sample processing and library preparation methodologies may confound interpretations based on viral abundance.

Here, we sought to parse the relative contributions of host and spatiotemporal factors to virus diversity and virome composition. To this end, we took a meta-transcriptomics approach to assemble a virome distribution dataset of rural and sylvatic mosquito species associated with vector activity in regions with history of arbovirus outbreaks. Specimens were repeatedly sampled from sites in French Guiana, Central African Republic, Madagascar, and Cambodia in a two-year sampling design to capture the temporal and seasonal stability of mosquito viromes. In addition, we examined the prevalence and abundance of certain viruses distinguished by their wide distributions, revealing high inter-species and inter-genus virome flux in Madagascar.

## RESULTS

### Virome characterisation of 18 mosquito species

Repeated mosquito samplings were performed during the dry and rainy seasons from late 2018 to early 2021 in French Guiana, Central African Republic, Madagascar, and Cambodia (Supplementary Figure 1). Mosquito species were initially identified based on morphological keys and later validated by the mitochondrial *cox1* DNA barcoding gene. We characterised the viromes of 18 mosquito species, including known disease vectors, from the genera *Aedes*, *Culex*, *Anopheles*, *Mansonia*, and *Coquillettidia* (Supplementary Table 1) through meta-transcriptomic sequencing on pools of five female mosquitoes of the same species, collection site, season, and collection period. Viral abundance and viral distribution matrices were generated for 16 of these species, excluding *Ae. serratus* and *Ma. africana* which has insufficient sample numbers. These matrices were later used in analyses to quantify the relative contributions of host and spatiotemporal factors to virus diversity and virome composition.

An average of 6.6 million reads were obtained per sequencing library. Following de novo assembly of reads, we removed libraries containing poor-quality contigs (*n* = 50). Libraries containing multiple host mosquito species (*n* = 23), identified though alignment against *cox1* gene sequences, were also removed. We detected 10,358 viral contigs (0.79% of total contigs above 1 kb) through shared homology with viral amino acid sequences recorded in the NCBI database. Viral contigs sharing >95% nucleotide sequence identity over >90% of contig lengths were clustered into 4,181 non-redundant viral operational taxonomic units (vOTUs). vOTUs were then identified by detection of specific viral hallmark genes: RNA-dependent RNA polymerase (RdRp) for RNA viruses, reverse transcriptase (RT) for reverse-transcribing viruses, DNA polymerase for double-stranded DNA viruses, replication-associated protein (Rep) for circular single-stranded DNA viruses, and NS1 for parvoviruses. Finally, 2290 contigs containing viral hallmark gene sequences representing 966 vOTUs were obtained from 381 sequencing libraries of *cox1*-validated mosquito species. To delineate novel viruses from those previously identified in the Pfam, ECOD, and NeoRdRp databases, we applied a <90% amino acid sequence identity threshold. Only 44 of these 966 vOTUs were known or previously reported (Supplementary Figure 2, Supplementary Table 2). It should be noted that this threshold is not universally applicable for all RNA virus lineages and we have selected a particularly conservative virus species demarcation threshold at the risk of overestimating virus diversity^22^.

From mosquito homogenates (*n* = 84) cultured in C6/36 *Ae. albopictus* cells for two blind passages, seven novel viruses were successfully propagated for the first time from *Ae. albopictus* and *Cq. venezuelensis* mosquitoes (Supplementary Table 2). The relatively low number of cultured viruses may reflect host ranges that exclude *Ae. albopictus* or the presence of viruses in our dataset that do not infect mosquitoes. Accordingly, we refer to all detected viruses as mosquito-associated, which encompasses both mosquito-infecting and non-mosquito-infecting viruses, to preserve the potentially meaningful biological links between viral and mosquito taxa from the same ecological context.

We performed a rarefaction analysis on virus species richness as a function of sequencing library numbers to visualise whether our number of sequencing libraries has captured the full virus diversity in each mosquito species. We opted for this level of analysis as our primary objective was to characterise the viral community composition at the species level, where individual sample pools represent replicates of the same biological unit rather than independent, heterogeneous environments. This approach ensures that our diversity estimates reflect the full virome diversity inherent to each host species, rather than being skewed by the inherent stochasticity of low-titre viruses in individual specimens. Rarefaction curves across 18 host mosquito species indicated that while discovered viral richness increased with sampling effort as expected, for most species, curves did not reach a plateau (Extended Data Figure 1a). This indicates that the current survey captured a significant portion of the species-specific virome, but there is more viral diversity to be discovered. Steep initial slopes, such as that observed in *Ma. uniformis*, suggests a relatively uneven distribution of individual viruses across sequencing libraries.

The 966 detected vOTUs belonged to 44 virus families, including single-stranded DNA, reverse-transcribing (RT), double-stranded, and positive and negative single-stranded viruses (Extended Data Figure 1b, Supplementary Figure 3). The distribution of these viral kingdoms varied considerably by species (Extended Data Figure 1b). *Mansonia* mosquitoes were almost entirely dominated by RT viruses, while *Culex* mosquitoes notably showed lower proportions of RT viruses relative to other mosquito genera. The prevalence of RT viruses across all mosquito species in our study suggests that retro-element-like sequences or specific reverse-transcribing virus families are a global core component of mosquito viromes.

Out of 2509 hallmark gene-encoding contigs, 219 (8.7%) were unable to be assigned to any ICTV-ratified virus family— where unclassified viral contigs make up between 0% to 28.6% of total viral contigs per mosquito species (Extended Data Figure 1c). The 0% value in *Cx. vishnui* samples is likely an artefact of low sample size and/or contig count, given its rarefaction curve. Amino acid sequence identities of contigs assigned to unclassified viruses mostly fell below 90%, indicating these are novel viruses (Extended Data Figure 1d).

### Diversity of mosquito viromes is largely determined by host species

Analyses on mosquito virus diversity and virome compositions were performed on a dataset containing only viruses detected in at least three sequencing libraries (*n* = 263 vOTUs). In addition, two mosquito species with exceptionally low sampling intensities, *Ae. serratus* and *Ma. africana*, were also excluded from downstream statistical analyses.

We measured virome alpha diversity using virus species richness values and Shannon index for each mosquito population, which we define here as an aggregated group of sequencing libraries representing the same mosquito species, collection period, site, and season (Figure 1a)^23^. For both measures, mosquito species is a significant predictor variable (Kruskal-Wallis, *p*-value < 0.05). Post-hoc pairwise comparisons revealed similarity groups in alpha diversity among mosquito species (Figure 1a) that do not appear to correlate with host phylogenetic relatedness. By both species richness (S_obs_) and Shannon index (H’), *Ma. titillans* (S_obs_ = 18; H’ = 1.92)*, Cq. venezuelensis* (S_obs_ = 8; H’ = 1.06), and *Ma. uniformis* (S_obs_ = 6; H’ = 1.07) harboured the highest virome diversity across all mosquito species examined.

**Figure 1.**
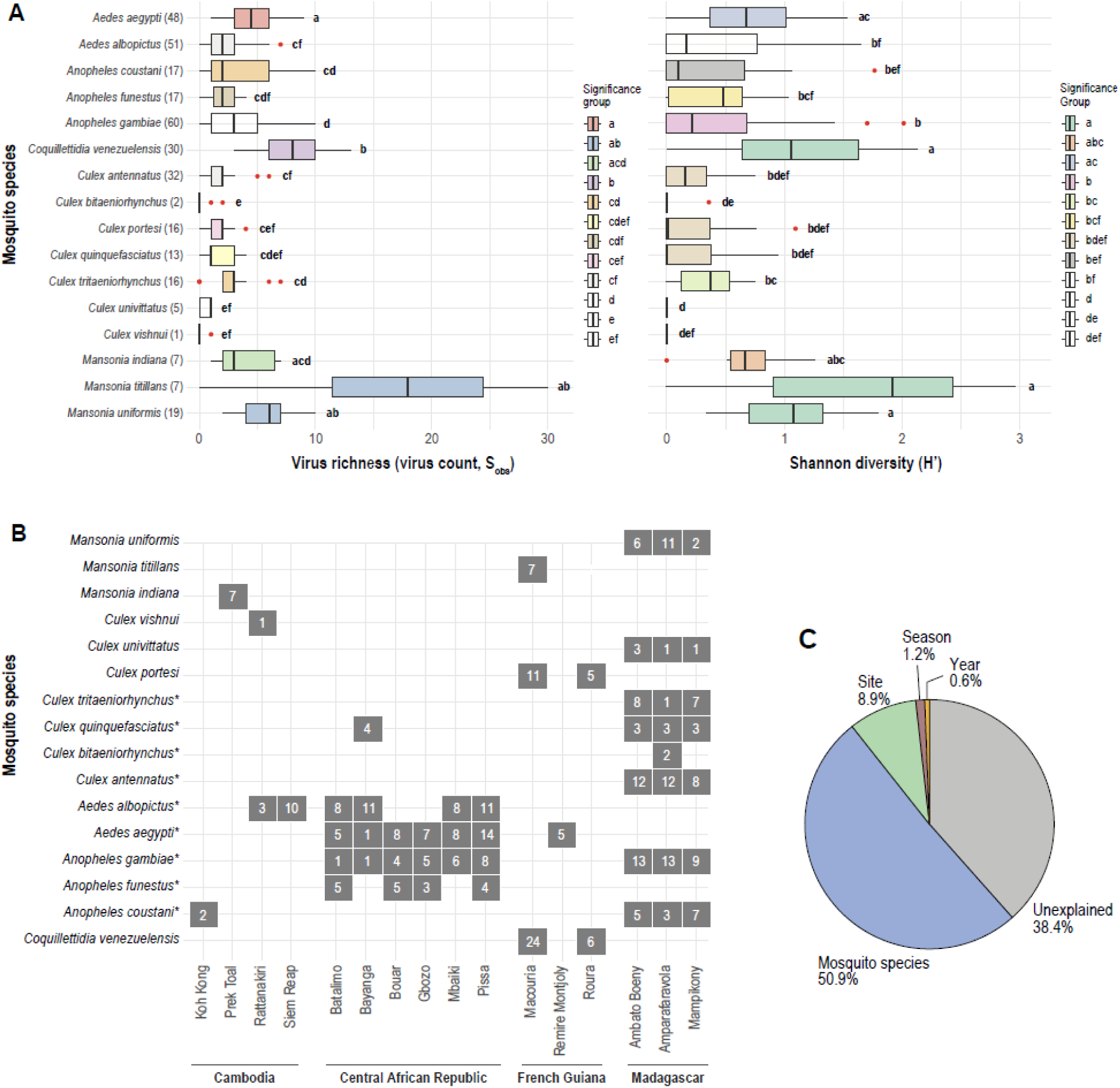
Factors driving virome diversity across mosquito populations. **a**, Virus species richness (S_obs_) and Shannon diversity indices (H’) for each mosquito species, displayed on a shared y-axis. Sample size (number of sequenced libraries) per mosquito species are indicated in brackets. Bar colours and letter codes denote statistical similarity groups determined by Dunn’s post-hoc pairwise comparisons following significant Kruskal-Wallis tests. **b,** Sample sizes or number of sequenced libraries for each mosquito species by site. Mosquito species sampled in more than one country are indicated by asterisks. **c,** Proportion of variance in mosquito virus community richness explained by individual factors.

To understand the influence and relative contribution of host and ecological factors on virome diversity, we fitted generalised linear models (GLM) for virus richness as a function of five possible predictors (mosquito species, collection site, season, and year). The best model, selected based on Akaike Information Criterion, included mosquito species, site, season, and year as predictors (Supplementary Table 3). By performing an analysis of deviance on this model, we found that these factors cumulatively explained 61.6% of variance (mosquito species: 50.9%, site: 8.9%, year 0.7%, and season: 1.2%) (Figure 1c), leaving 38.4% unexplained.

### Mosquito virome composition is driven by host and spatial factors in a virus-dependent manner

A principal coordinates analysis (PCoA) on a Jaccard distance matrix computed from aggregated virus species presence-absence data across mosquito populations revealed strong influence of host phylogeny (Extended Data Figure 2). Populations appear to cluster according to host genus independent of country or season. A *Coquillettidia* cluster was observed isolated from other mosquito genera, while *Culex* mosquitoes displayed the broadest virus distribution across the ordination space. The top two axis explained 5.2% and 4.5% of total variance, respectively. These low values indicate that virome composition is under the influence of many factors with relatively small effect sizes. Tests for multivariate dispersion showed stable within-group virome variance across locations and seasons, validating the PERMANOVA results (Supplementary Table 4). Mosquito communities in Madagascar, Cambodia, and Guyane shared nearly identical compositional spread without outlier groups, and seasonal variation was minimal. Overall, these results confirm that significant community differences reflect true compositional shifts rather than artifacts of unequal dispersion.

To assess how individual spatiotemporal factors structure virome composition within each mosquito species, we then performed a permutational multivariate analysis of variance (PERMANOVA)^24^ on the Jaccard distance matrix (Table 1). R^2^ values represent the proportion of total variation in virome composition explained by each term. Across the 16 mosquito species, site or country explained a significant proportion of variation (*p*-value < 0.05) in all species except *Cx. vishnui*. For these this species, we likely had limited statistical power to detect site effects due to its small sample size. By contrast, season was a significant variable only for three species, and in each case the variation attributable to site was much more substantial than that attributable to season. Together, these results indicate that mosquito virome composition is structured primarily by spatial factors rather than by temporal factors.

**Table 1.**
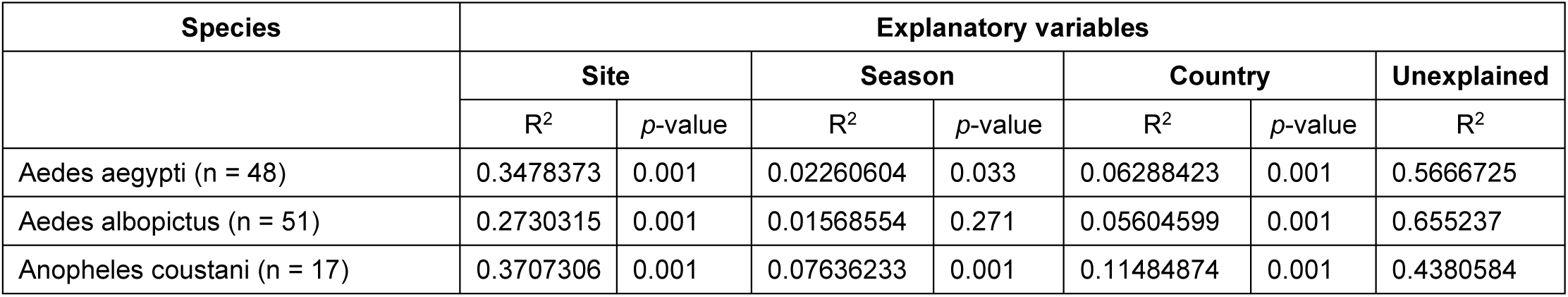
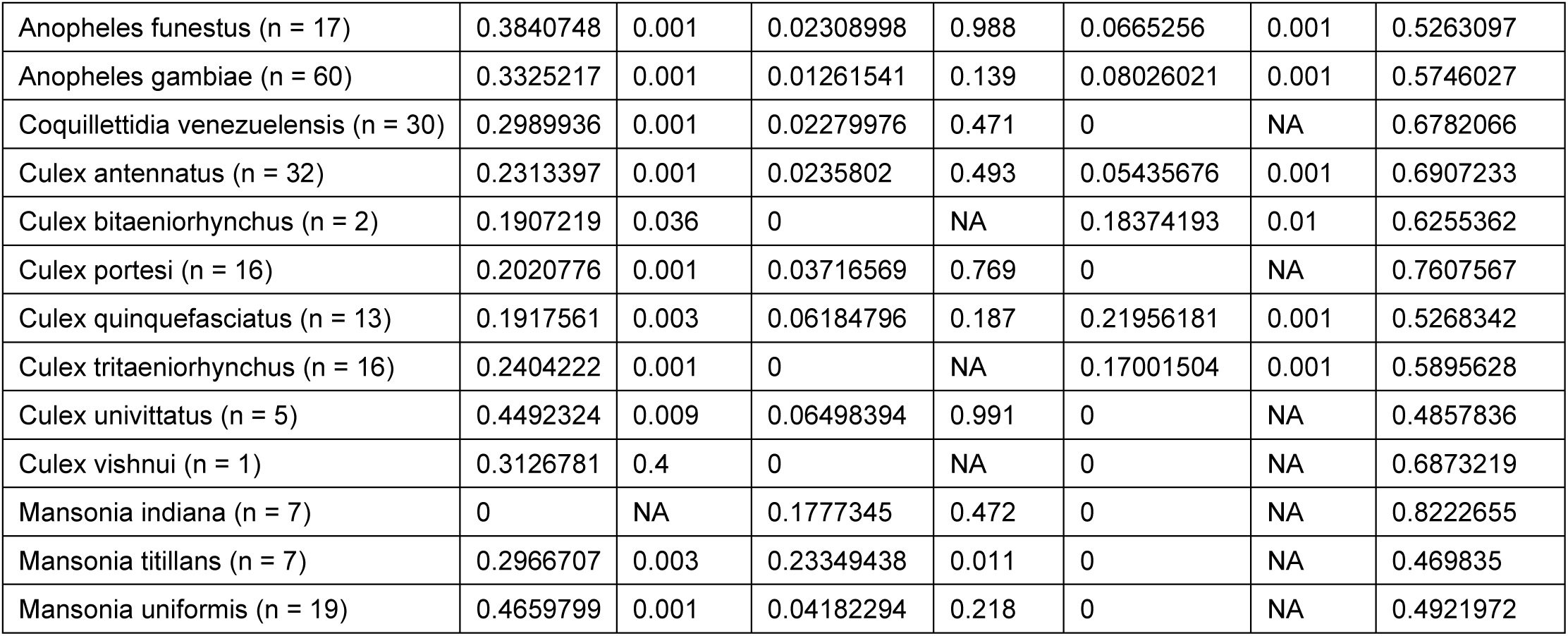
Output of hierarchical permutational multivariate analysis of variance (PERMANOVA) on a Jaccard distance matrix derived from viral species presence-absence data. Only mosquito species represented by at least three sequencing libraries and viruses detected in at least three individual sequencing libraries were included in the analysis. R-squared (R²) values above 0.20 indicate strong relative contribution of a given predictor variable predictive effect, whereas values lower than 0.05 indicate weak relative contribution. Significant *p*-values (< 0.05) are shown in bold.

We next performed an indicator species analysis on viral composition data to reveal significant associations between individual viruses with a particular host (mosquito genus or species), spatial (country or site) and temporal (season) variable. Significant viral indicators (*p*-value < 0.05) are visualised in a consolidated dot plot (Figure 2). The presence of a dot indicates a significant association, while its size, representing the indicator value index, is a measure of a virus’s specificity and fidelity to a particular condition or category within a variable. Across analysed virus taxa, host and spatial factors were the strongest predictors of a virus presence within a given community (Table 2). We reveal host specialist viral taxa that are strongly associated with specific host or ecological context contributing to the distinct virome compositions observed in the PERMANOVA. For example, *Phasivirus phasiense* (Phasi Charoen-like phasivirus) is widely reported across previous virome studies in *Ae. aegypti* mosquitoes. In our dataset, this virus is highlighted as an indicator for this mosquito species (Figure 2), in which it is the most abundant virus (Extended Data Figure 3).

**Figure 2.**
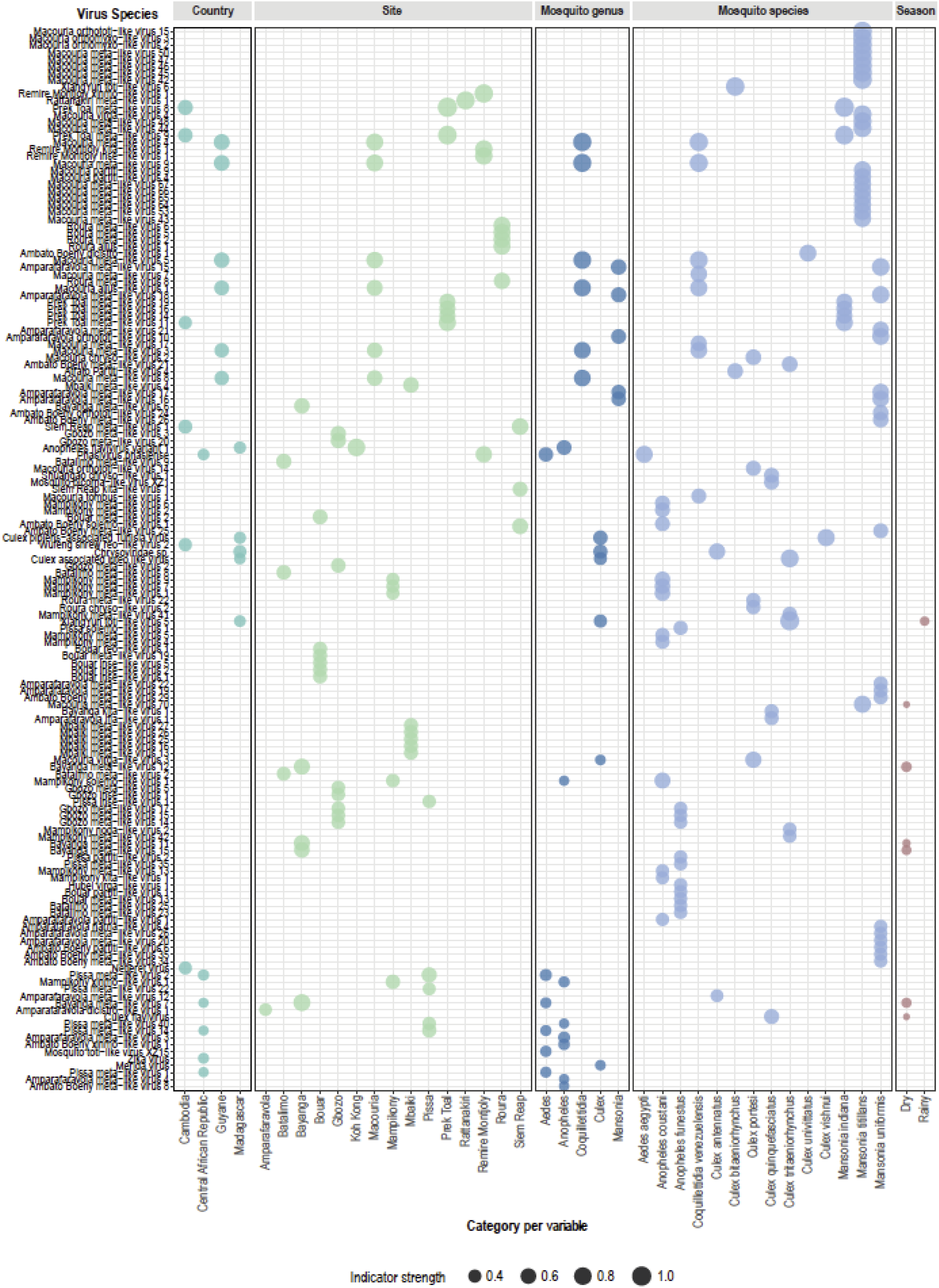
Indicative associations between individual viruses and host or ecological indicators. A dot plot depicting the top significant indicators (*p*-value < 0.05) of virus species (y-axis) ranked by their indicator value strength. Dot size represents the specificity and fidelity of the virus to a particular host or ecological context. Only significant associations are shown.

**Table 2.** Twenty viruses with the highest significant indicator value (IndVal) index.

| Virus | Predictor term | Associated variable | IndVal index | p-value |
| --- | --- | --- | --- | --- |
| Prek Toal meta-like virus 8 | Mosquito species | *Mansonia indiana* | 1 | 0.001 |
| Prek Toal meta-like virus 8 | Site | *Prek Toal* | 1 | 0.001 |
| XiangYun toti-like virus 5 | Mosquito species | *Culex tritaeniorhynchus* | 0.968 | 0.001 |
| Prek Toal meta-like virus 9 | Mosquito species | *Mansonia indiana* | 0.926 | 0.008 |
| Macouria orthototi-like virus 15 | Mosquito species | *Mansonia titillans* | 0.926 | 0.004 |
| Macouria meta-like virus 42 | Mosquito species | *Mansonia titillans* | 0.926 | 0.004 |
| Macouria orthomyxo-like virus 3 | Mosquito species | *Mansonia titillans* | 0.926 | 0.004 |
| Macouria orthomyxo-like virus 2 | Mosquito species | *Mansonia titillans* | 0.926 | 0.004 |
| Macouria meta-like virus 47 | Mosquito species | *Mansonia titillans* | 0.926 | 0.004 |
| Macouria meta-like virus 45 | Mosquito species | *Mansonia titillans* | 0.926 | 0.004 |
| Macouria meta-like virus 46 | Mosquito species | *Mansonia titillans* | 0.926 | 0.004 |
| Macouria meta-like virus 50 | Mosquito species | *Mansonia titillans* | 0.926 | 0.004 |
| Prek Toal meta-like virus 9 | Site | *Prek Toal* | 0.926 | 0.001 |
| XiangYun toti-like virus 6 | Mosquito species | *Culex bitaeniorhynchus* | 0.913 | 0.007 |
| Remire Montjoly xinmo-like virus 1 | Site | *Remire Montjoly* | 0.894 | 0.002 |
| Macouria meta-like virus 4 | Mosquito species | *Coquillettidia venezuelensis* | 0.876 | 0.001 |
| Macouria meta-like virus 4 | Mosquito genus | *Coquillettidia* | 0.876 | 0.001 |
| Rattanakiri meta-like virus 1 | Site | *Rattanakiri* | 0.866 | 0.001 |
| Macouria meta-like virus 9 | Mosquito species | *Coquillettidia venezuelensis* | 0.856 | 0.001 |
| Macouria meta-like virus 9 | Mosquito genus | *Coquillettidia* | 0.856 | 0.001 |

### Virus sharing across mosquito populations

To understand how host and ecological factors shape virus sharing across spatially and temporally distinct mosquito populations, we constructed a virome similarity network based on Jaccard similarity coefficients (Figure 3). Each node represents a group of sequencing libraries from the same mosquito species, site, and collection period (season and year). Across the network, multiple small, connected clusters were formed from two to five nodes representing the same mosquito species. Apart from one cluster of *Ae. albopictus* populations from Siem Reap and Rattanakiri in Cambodia, all single mosquito species clusters also originated from the same site. This suggests that site-level fidelity remains a powerful determinant of virome structure within a given mosquito population alongside host taxonomy.

**Figure 3.**
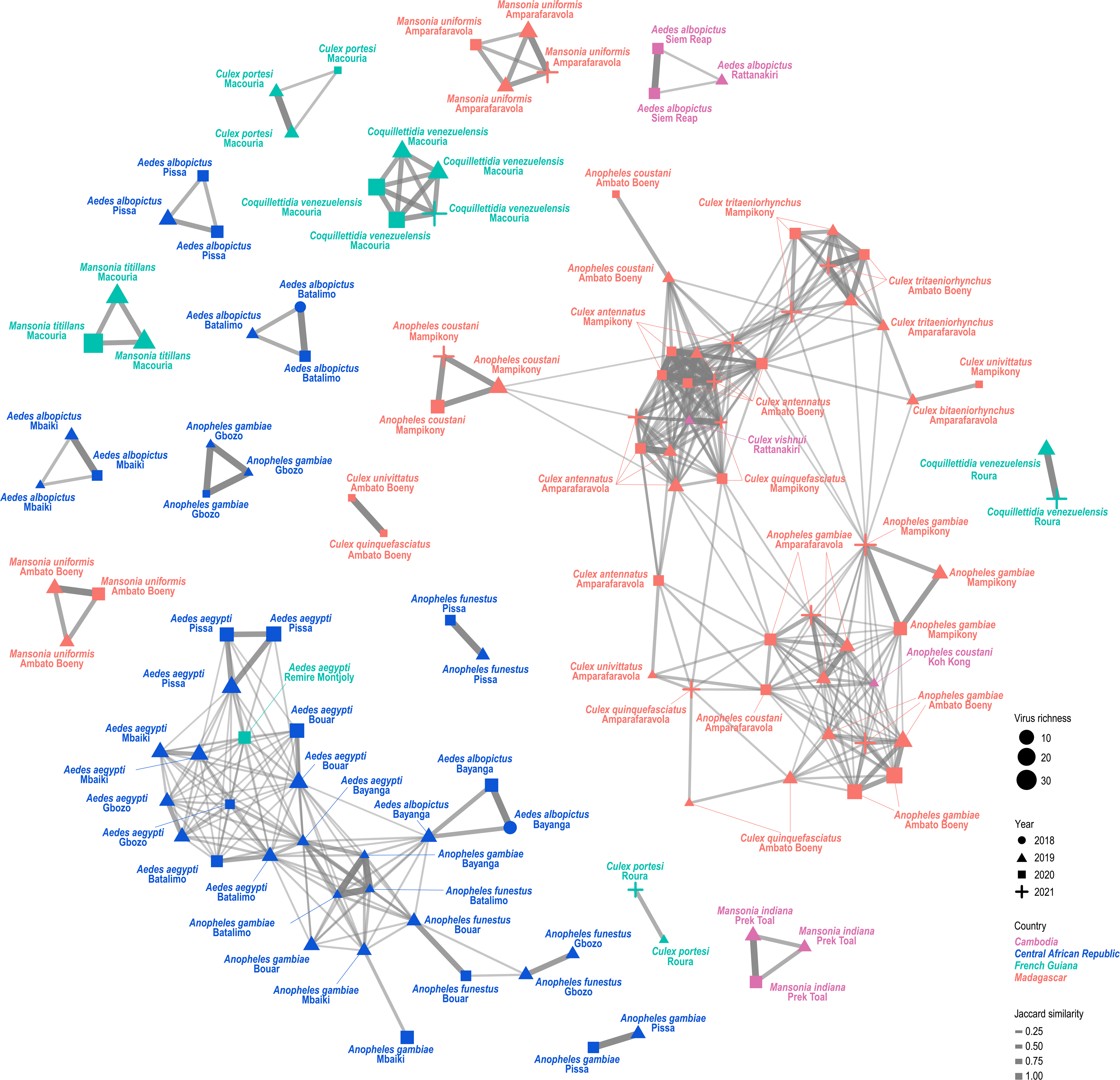
High frequency of virus sharing among African mosquito populations. Virus similarity network in a Fruchterman–Reingold layout, constructed from Jaccard similarity coefficient matrix. Each node represents a group of sequencing libraries from the same mosquito species, site, and collection period (season and year). Edges represent shared virus species between two nodes, the thickness indicates the number of shared viruses. Only edges with Jaccard similarity above 0.05 are shown.

Two large mixed-species clusters emerged, distinguished spatially at the country level (Figure 3). A cluster of predominantly Madagascar populations included *Culex* and *Anopheles* species, while a second cluster was composed of *Aedes* and *Anopheles* species from the Central African Republic. Within each of these major clusters, Jaccard similarity values still tended to be higher between conspecific nodes. Notably, although *An. gambiae* were present in both the Central African Republic and Madagascar, Jaccard similarity index indicated low or no similarity between the two geographically separated populations. No segregation corresponding to year was observed.

### Using Madagascar as a model to examine regional virome dynamics

The insular biogeography of Madagascar and the particularly even sampling performed across sites, seasons, and years provided an ideal country-level subset of virome data to examine virus-host associations more closely (Figure 1B, Supplementary Figure 1a). Madagascar is the largest contributor to the global virome diversity of this study, being the origin of 80 viruses out of the total of 263 (post *n* > 3 filter). Across the three sites, the diversity of mosquito species sampled and the number of virus species detected per sequencing library were comparable (Figure 4a-b). Although Mampikony was represented by slightly fewer sequencing libraries, this did not appear to be reflected in the number of detected viruses.

**Figure 4.**
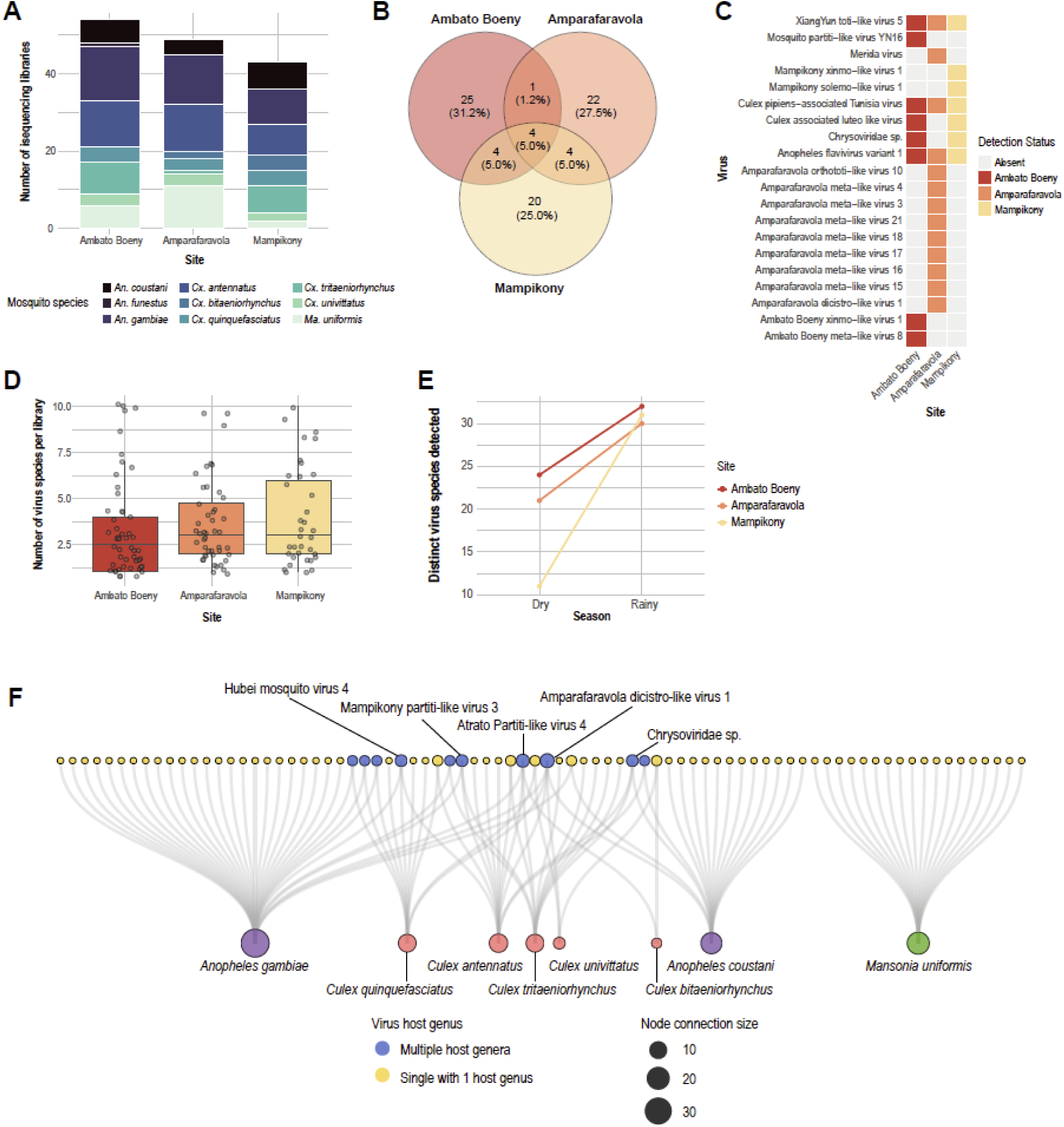
The virome structure of Madagascar. **a**, Proportion of mosquito species sampled per site. **b,** Number of shared and site-specific viruses. **c,** Distribution of the top 20 most abundant viruses across sites. **d,** Virus species richness across three sites. Each data point represents a sequencing library. **e,** Virus species richness by site across seasons. **f,** Bipartite network showing virus–host associations. Virus node colours denote host specialism or generalism (associated with more than one mosquito genus).

Mampikony shared eight viruses each with Ambato Boeny and with Amparafaravola (Figure 4b). This may be explained by its geographical location, which sits between Ambato Boeny to the north and Amparafaravola to the south (Supplementary Figure 1b). This suggests the interactive influence of ecology (spatial distance) and evolutionary (host specificity) factors on virus distribution across spatial gradients.

Virome alpha diversity, measured by viral richness values, were lower than expected given the number of viruses detected in each site (Figure 4b, d). This suggests that sampled mosquitoes harboured small but highly variable virus communities. That is, individual viruses are sparsely distributed across a given mosquito population at low prevalence. Although we observed a clear increase in the number of virus species found during the rainy compared to the dry seasons across all sites (Figure 4e), this could not be decoupled from the effects of less intensive sampling during drier climate due to reduction in mosquito population size.

Four viruses, Anopheles flavivirus variant 1, Culex pipiens-associated Tunisia virus, XiangYun toti-like virus 5, and Atrato Partiti-like virus 4, were detected in every site, suggesting substantial virome flux between spatially separated populations (Figure 4b-c). Collectively, viruses were associated with *An. gambiae*, *Cx. antennatus*, *Cx. bitaeniorhynchus* and/or *Cx. tritaeniorhynchus* mosquitoes. The common presence of host mosquito species in every site explained the distributions of three out of the four viruses (Extended Data Figure 4).

To identify generalist or specialist viral host ranges in the Madagascar context, we constructed a bipartite network between viruses and their associated host mosquito species. Each mosquito species was associated with at least two viruses. Fifteen viruses were associated with more than one host species. Among these, ten exhibited cross-generic host ranges. Specifically, these viruses were shared between *Anopheles* and *Culex* mosquitoes (Figure 4f). Notably, Atrato partiti-like virus 4 was associated with four different host species, giving rise to its broad distribution (Extended Data Figure 3, Figure 4f). However, this virus displayed relatively low abundance compared to other viruses associated with its hosts (Extended Data Figure 4).

### Viruses with broad host ranges exhibit varying abundances

We next investigated virus-host associations at a broader spatial scale using our global dataset. To focus on more biologically meaningful associations in the absence of experimental infection data, that is, those that are more likely to reflect true infections, we limited our analysis to the 50 most abundant viruses. The mean normalised viral read abundance within this subset ranged from 4,661.94 to 339,422.40 reads per million (RPM).

Few viruses displayed broad spatial ranges, which could often be attributed to their association with metropolitan host mosquito species. *Phasivirus phasiense* (Phasi Charoen-like phasivirus) and Mosquito toti-like virus XZ15 were detected in the Central African Republic as well as French Guiana, where their single host species, *Ae. aegypti*, originated from (Figure 5a, Extended Data Figure 4). Similarly, Anopheles flavivirus variant 1 in Koh Kong, Cambodia, was associated with *An. coustani*, which was also sampled in Madagascar (Figure 5a, Extended Data Figure 4). However, Culex pipiens-associated Tunisia virus in Cambodia was detected in *Cx. vishnui*, a species specific to the region. In Madagascar, this virus was detected instead in *Cx. antennatus* and *Cx. tritaeniorhynchus* (Figure 5a, Extended Data Figure 4), demonstrating an example where host generalism contributes to broader spatial ranges and geographical prevalence.

**Figure 5.**
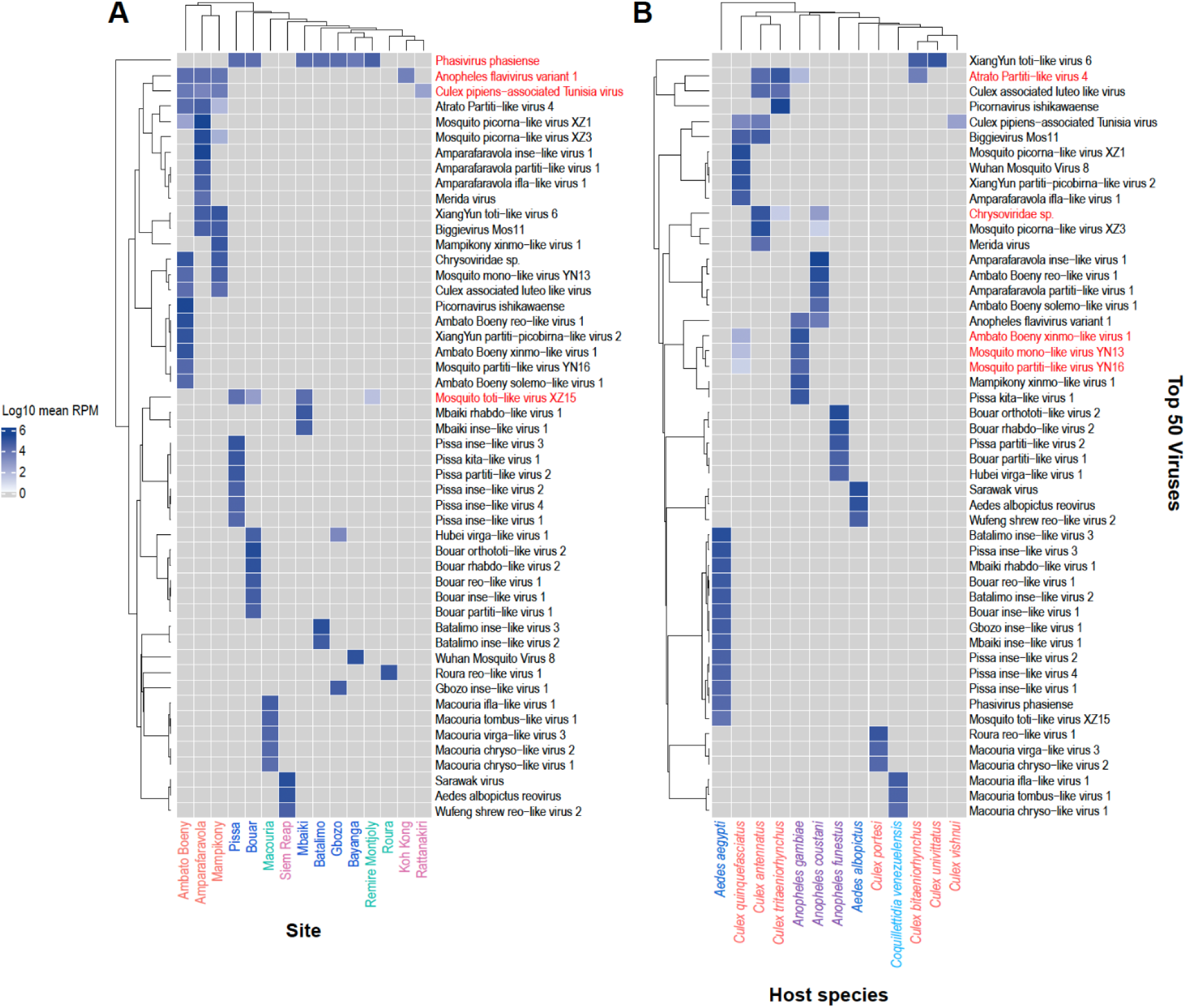
Distribution of the most abundant viruses across host species and site. Heatmap depicting the abundance, measured by mean normalised reads per million (RPM), as an indication of likelihood of true infection of the top 50 abundant viruses (y-axis) across **a,** site and **b,** host species. X-axes label colours denote **a,** country (Cambodia, pink; Central African Republic, blue; French Guiana, green; Madagascar, orange) or **b,** host genus. Virus names in red indicate those that are shared **a,** across countries and **b,** across host genera.

The cross-genus virus sharing patterns observed among Madagascar mosquitoes are also reflected here; Five viruses, including Atrato Partiti-like virus 4, Chrysoviridae sp., and Ambato Boeny Xinmo-like virus 1, are associated with *Culex* and *Anopheles* mosquitoes (Figure 5a-b). Notably, all cases of within-genus virus sharing were also only observed among *Culex* or *Anopheles* species.

Viral abundance data revealed further nuance into the infection dynamics of viruses with broad host ranges. Within-genus shared viruses, such as XiangYun toti-like virus 6 or Culex pipiens-associated Tunisia virus, exhibited similar abundance levels across their different host species (Figure 5a). By contrast, cross-genus shared viruses showed greater variation in viral abundance across hosts, displaying a bias towards one particular host species or genus. Specifically, Atrato Partiti-like virus 4 was present in high abundances in its three *Culex* hosts relative to *An. gambiae*. Similarly, Chrysoviridae sp. was detected more abundantly in *Cx. antennatus* compared to *Cx. tritaeniorhynchus* and *An. coustani*. Ambato Boeny xinmo-like virus 1, Mosquito mono-like virus YN13, and Mosquito partiti-like virus YN16 were more abundant in *An. gambiae* than in *Cx. quinquefasciatus*. These variations in viral abundances may reflect the evolutionary histories of host range expansion of these viruses.

## DISCUSSION

We investigated the virus communities of 16 mosquito species in four countries across a two-year period to understand the influence of host and ecological factors in shaping virome dynamics. Host species emerges as the strongest determinant of virome richness or virus distribution in mosquito populations, in line with previous studies^2,15,16^. Season and collection year have limited effects on virome diversity and composition, suggesting relatively stable virome composition over time. Individual virus species are differentially influenced by host phylogeny or spatial factors, highlighting the importance of specific virus-host relationships underlying virome distribution.

What other factors could be driving virome dynamics? Many more unexamined variables may explain the residual variance in virus diversity and virome composition. Spatiotemporal variation is likely tied to the diversity of plants, ecosystem disturbance level, microclimatic patterns, or mosquito species richness within a specific ecosystem. A previous study suggested that these factors, in particular the latter two, also act in a heterogenous and virus-specific manner^25^. While we have focused on female mosquitoes, a recent study showed that within-species virome diversity varied by mosquito sex, presumably linked to the hematophagous behaviour of female mosquitoes^26^. Viral communities in female mosquitoes are likely perturbed by post-bloodmeal immune stimulation and metabolic changes^27,28^. The presence of virus-transmitting ectoparasites of mosquitoes, such as water mites^29^, should also be examined as a potential factor shaping virome composition, as observed in a study in honeybees^30^.

Examining the environmental viromes of mosquito habitats, especially aquatic larval habitats, would invaluably shed light on the potential transmission pathways of non-vertebrate-infecting mosquito viruses. In this study, rather than distinguishing true mosquito-infecting viruses from those of other mosquito-associated microorganisms (bacteria, fungi, parasites), we took a more holistic view of mosquito-associated viruses. These associations can provide important information on the ecological context of mosquito specimens, which can be revisited as we gain a deeper understanding of mosquito virome biology.

Our dataset also revealed high virome connectivity across mosquito populations in Madagascar. This could stem from a combination of evolutionary (host susceptibility factors or viral phenotypes supporting broad host ranges) and ecological (insular habitat biodiversity) factors. This uncovers several potential lines of further investigation into the evolutionary history of specialist and generalist viruses. We define the latter as viruses exhibiting broad host ranges spanning two mosquito genera. We observed incidents of virus sharing among *Culex* species in Madagascar and Cambodia, as reported by earlier studies in Senegal^17^ and Belgium^31^. Interestingly, cross-genus virus sharing occurred most frequently between *Culex* and *Anopheles.* As not all five mosquito genera were evenly represented in our dataset, i.e., few *Culex* specimens were collected in the Central African Republic and no *Aedes* specimens were obtained from Madagascar, we cannot draw definitive conclusions on the lack of virus sharing between *Aedes* and *Culex* hosts. Similarly, an important caveat to reports of species-specific viruses is the possibility that a divergent, co-occurring host species was unsampled.

Caution is warranted as to whether mosquitoes are the true host of generalist viruses. Among the five viruses shared between *Culex* and *Anopheles* mosquitoes in Madagascar, three are closely related to the *Partitiviridae* and *Chrysoviridae* families. Although these viral families are historically associated with plant and fungi host, closely related viruses are often detected in mosquito virome studies^4^, potentially forming previously unknown lineages. Further evidence from infection experiments is therefore needed to validate these viruses as part of the mosquito infectome and to enable accurate classification of discovered mosquito viruses as host specialists or generalists.

Consistent with previous studies of these species from other biogeographic regions^13,32^, our dataset also revealed species- and genus-specific virome patterns in *Ae. aegypti* and *Ae. albopictus*. Of note, Phasi Charoen-like phasivirus (*Phasivirus phasiense*) is almost exclusive to *Ae. aegypti* despite its remarkably high prevalence across global populations^4,13,33^. This host specificity contrasts with what is known about *Aedes*-transmitted arboviruses (dengue, Zika, chikungunya, yellow fever viruses)^34^, which uses principally *Ae. aegypti* and *Ae. albopictus* as vectors but may also be transmitted by sylvatic *Aedes* species such as *Ae. malayensis*, *Ae. scutellaris, or Ae. vittatus*^35–38^. Notably, *Culex*-associated arboviruses (West Nile and Usutu viruses) have multiple reported vector species^34,39^. This provokes the question: Are mosquito viruses with broad host ranges more likely to host-jump to vertebrates and emerge as novel pathogens? Addressing this question through deeper virome surveillance would allow the identification of potential novel arboviruses with high-risk of emergence.

*Culex* mosquitoes exhibited particularly rich viromes, yet it remains unclear how virome richness correlates with vectorial capacity. Through dilution effects in within-host pathogen dynamics^40^, high virome diversity may reduce the frequency of hosts with hospitable viromes for arboviruses to invade and establish infections in^41^. This may be underpinned by mechanisms such as superinfection exclusion or competition for shared metabolic resources. Conversely, low diversity viromes may reflect refractory host factors or robust antiviral immunity. Resolving this dichotomy will require targeted investigation into correlations between within-species variation in virome diversity and mosquito basal immunity.

Mosquito viromes may directly influence their vectorial capacity through interactions with co-infecting arboviruses^6^. Several mosquito-specific viruses have been documented to inhibit co-infecting arboviruses^9,10,12,14,42^, making them a promising biological tool to reduce vector competence^6,43^. Yet, virome-based biocontrol strategies via the release of mosquitoes harbouring anti-arboviral viromes fundamentally require a solid understanding of which ecological factors shape virome composition and how stable viromes are over time. Our finding that mosquito viromes are relatively invariable to temporal factors is encouraging, as it implies that an engineered virome can sustainably persist in natural populations over time. However, predicting how an introduced virus will spread across an ecosystem will require careful consideration of specific host and viral taxa combinations, supported with robust experimental evidence.

Whether through surveillance or biocontrol measures, elucidating virome dynamics can lead to improved pre-emption and management of arbovirus outbreaks. Our study revealed deep complexities in the interplay between evolutionary and ecological drivers of mosquito virus diversity and virome composition, uncovering new hypotheses-driven questions about virus-mosquito interactions in their natural ecologies in the context of emergence of novel pathogens.

## METHODS

### Mosquito sample collection and processing

Collections of adult mosquitoes were undertaken in French Guiana, Central African Republic, Madagascar, and Cambodia from November 2018 to March 2021 during dry and rainy seasons. These four countries lie within the tropical region (latitudinal bounds 23.5° north and south of the equator). This study includes 16 collection sites, which comprised communes, villages, and towns in French Guiana (n = 3), Central African Republic (n = 6), and Madagascar (n = 3), and national reserves in Cambodia (n = 4) (Supplementary Figure 1). These sites represent five distinct bioregions as defined by the One Earth Bioregions Framework^44^ (Guianan Forests and Savannahs, Mandara Mountain and North Congolian Forest-Savannahs, North Congolian Lowland Forests, Madagascar Island, and Indochina Mixed Forests & Peatlands). All sites were classified as the Tropical & Subtropical Moist Broadleaf Forests biome, with the exception of five sites in the Central African Republic, which belonged to the Tropical & Subtropical Grasslands, Savannas & Shrublands biome^45^ (Supplementary Table 1).

A combination of trapping methods including CDC light traps, CO_2_-baited BG sentinel traps, and human-landing catches were used. Initial species identification was performed in-field by medical entomologists using geographically relevant morphology keys^46–56^ on cold tables before preservation in liquid nitrogen and transport on dry ice to laboratories. In addition, mosquito species identity was later bioinformatically confirmed using *cox1* DNA barcodes as described below. Mosquito specimen information is provided in Supplementary Table 1.

Female mosquito specimens of the same species, sampling site, and collection period (season and year) were grouped into pools of five individuals. To prepare samples for RNA extraction and virus isolation, samples were homogenized with a Precellys (Bertin Instruments) at a speed of 6000 rpm for 30 seconds in 500 μL of non-supplemented Leibovitz’s L-15 medium (Gibco, 11415049). Mosquito homogenates were clarified by two rounds of centrifugation at 7500 xg for 15 minutes at 4 °C to remove debris.

### RNA isolation, library preparation and sequencing

Total RNA was isolated from 200 μL of clarified homogenate using 300 μL of TRIzol reagent (Invitrogen, 15596026) as per the manufacturer’s protocol. Quantification of RNA was performed using a Qubit RNA High Sensitivity Assay Kit (Invitrogen, Q32855) on a Qubit Fluorometer (Invitrogen). cDNA libraries were prepared using the Zymo-Seq RiboFree Total RNA Library Kit (Zymo Research, R3003), which universally removes stoichiometrically abundant host ribosomal RNA (rRNA). Meta-transcriptomic (total RNA) sequencing of 528 dual-indexed libraries was performed on a NextSeq 500 system (Illumina), generating ∼3 billion 150 bp single-end reads, obtaining a sequencing depth of >5 million reads per library. Sequencing library quality was assessed using FastQC, and adapters were removed using Fastp 1.0.1.

### Contig assembly and virus detection

Raw reads from each library were de novo assembled using *SPAdes* 4.2.0 under rnaviralSPAdes parameters. Libraries containing contigs of poor assembly quality were removed from further analysis (*n* = 50). To detect viral contigs, we performed a homology search on contigs more than 1 kb in length with *DIAMOND blastx* option under very-sensitive parameters (E-value cutoff at 1 ⋅ 10^-5^) against the NR non-redundant protein database from NCBI. A total of 10,358 contigs homologous to known viral sequences were identified (0.79% of total number of contigs >1 kb).

Viral contigs that share >95% nucleotide sequence identity over >90% of contig lengths were clustered into non-redundant sequence clusters using *BLASTn*. This high identity threshold was selected to enable the detection of distinct strains of the same virus species within our study. A total of 4,181 viral operational taxonomic units (vOTUs) were identified. Their identity was confirmed by detecting hallmark genes specific to each viral group: RNA-dependent RNA polymerase (RdRp) for RNA viruses, reverse transcriptase (RT) for reverse-transcribing viruses, DNA polymerase for double-stranded DNA viruses, replication-associated protein (Rep) for circular single-stranded DNA viruses, and NS1 for parvoviruses. These hallmark genes were identified by searching translated viral contigs (Prodigal v2.6.3) against the Pfam, ECOD, and NeoRdRp databases using HHSearch or HMMER. 2,290 contigs containing viral hallmark genes, representing 966 viruses, were used for downstream bioinformatics analyses (Supplementary Table 2).

To identify novel species, these contigs were compared against viral sequences from two landmark invertebrate studies ^2,57^ and the NCBI NR database using *DIAMOND blastx*. We defined novel viral species using thresholds of <80% nucleotide identity and <90% protein identity.

### Viral read abundance quantification

To quantify viral abundance, raw reads were mapped against a reference set of viral contigs using bowtie2 in ‘--very-sensitive-local’ mode. These counts were subsequently normalised to Reads Per Million (RPM) by dividing the number of mapped reads for each contig by the total library read count and multiplying by 10^6^.

### Validation of host mosquito species using *cox1* gene sequence

Host mosquito species was validated for each sequencing library by mapping contigs containing the mitochondrial cytochrome *c* oxidase I (*cox1*) gene against the BOLD database^58^. *Cx. vishnui* here refers to the collective subgroup as individual species cannot be distinguished by the *cox1* marker alone. Libraries containing mixed mosquito species were removed (*n* = 23).

### Virus isolation

Clarified mosquito homogenates were cultured in *Aedes albopictus* C6/36 cells (ATCC, CRL-1660) for two blind passages. These cells are maintained at 28 °C in Leibovitz’s L-15 medium supplemented with 1% (v/v) of Penicillin-Streptomycin (Gibco, 15140122), 1% MEM Non-essential amino acid solution (Sigma, M7145), 2% Tryptose Phosphate Broth (Gibco, 18050039), with 10% heat-inactivated foetal bovine serum (Gibco, A5256701). To prepare for inoculation, confluent monolayers of cells in 24-well cell culture plates were washed with 500 μL of 1X Dulbecco’s phosphate-buffered saline (Gibco, 14190094). Cells were then inoculated with 50 μL of mosquito homogenate diluted in 150 μL of serum-free L-15 medium and allowed to incubate for one hour at 28 °C. Virus inoculum was then removed from each well and replaced with 1 mL of L-15 medium with 2% of serum and 2% of Amphothericin B (Eurobio Scientific, CABAMB00-0U); all other supplements as described above. Viruses were allowed to propagate in cells for seven days at 28 °C and cells were monitored daily for the presence of cytopathic effect. For subsequent passage, 200 μL of culture medium was inoculated onto fresh cells in 24-well cell culture plates, with inoculum removal and culture medium replacement as described above. At seven days post-inoculation, P_2_ virus isolates were collected and tested by virus-specific RT-PCR.

### Mosquito and virus phylogeny analyses

Viral contigs encoding hallmark genes RdRp, RT, DNA pol, Rep, and NS1 were used to perform a maximum likelihood phylogenetic analysis to determine virus taxonomy. Virus trees were constructed based on previous mosquito virome studies^2,57^ and were built at the super-clade level using hallmark genes. The trees were aligned using *MAFFT*-AUTO method^59^, trimmed using *trimAl*^60^, and were generated with *IQ-TREE 2* using the options - m MFP+C20 -B 1000 -alrt 1000 -nstop 500 to specify automatic best model selection (-m), perform ultrafast bootstrap (UFBoot) with 1000 replicates (-bb), and SH-like approximate likelihood ratio test on 1000 replicates (-alrt)^61,62^. UFBoot estimates branch support values with shorter computation time compared to standard bootstrapping^63^.

The mosquito species maximum-likelihood phylogenetic tree was constructed based on the mitochondrial *cox1* gene^58^. 596-bp long sequences were aligned using *MAFFT*-AUTO and was inferred using *IQ-TREE 2* using the same options stated above. The tree was rooted to water mite *cox1* sequences (Accession numbers MH916807, MN362807, KM101004).

## Statistical analysis

Statistical analyses on the diversity of mosquito viromes were performed at the virus species level on 16 mosquito species. *Ae. serratus* and *Ma. africana* libraries were excluded due to exceptionally low numbers. Only viruses detected in at least three individual sequencing libraries were retained for analyses. Rare singleton virus species that are only found in one sample pool may artificially inflate virus diversity estimates of mosquito populations and confound comparative analyses on virome composition.

### Rarefaction analysis of virus species diversity

Rarefaction measures and compares taxonomic diversity in datasets adjusting for sampling size^64^. Rarefaction analysis was performed to determine whether viral richness was sufficiently represented within each host mosquito species. A rarefaction curve was estimated for each mosquito species by repeatedly subsampling individual sequencing libraries (up to the total number of pools available for that species) and plotted against the average number of unique viral taxa found over 50 iterations. This gives the expected number of virus species to be found at a given number of sequencing library.

### Species biogeography illustration

Geographic coordinates for each collection site were obtained using the Google Maps Geocoding API through the *ggmap* package in R. Sites were geocoded by combining site names with their respective countries to ensure accurate geographic placement. For each collection site, mosquito species counts were calculated and summarized and shown in a pie chart. Base map data were obtained from Natural Earth using the *rnaturalearth* package, providing country boundaries at medium resolution.

### Alpha diversity metrics calculation

To measure the diversity of mosquito viral communities, two complementary alpha diversity metrics were calculated for each mosquito species. Data were first aggregated by summing Reads Per Million (RPM) for each viral species per sample pool. Viral species richness (S_obs_) was calculated as the total count of these filtered viral species present per sample. The Shannon diversity index (H′) was calculated using the *diversity()* function from the *vegan* package, utilizing the formula H′=−∑(pi×ln(pi)), where pi represents the proportional abundance of each viral species based on RPM values.

Non-parametric Kruskal-Wallis tests were performed with mosquito species as the primary predictor variable for both viral richness and Shannon diversity. Post-hoc pairwise comparisons among mosquito species were conducted using Dunn’s tests with Benjamini-Hochberg (BH) false discovery rate correction via the *rstatix* package. Compact Letter Displays (CLD) derived from Dunn’s adjusted p-values (*multcompView* package) were visualized using boxplots and bar plots; species sharing letters and colours do not differ significantly (p-value > 0.05).

### Generalised linear modelling

To investigate the drivers of alpha diversity in mosquito virus communities, virus richness was modelled using penalised generalised linear models (GLM) with Poisson error distribution and log-link function utilizing the *brglm2* package (brglmFit method)^65^. The validity of the Poisson error distribution was assessed by testing for residual overdispersion using the *dispersiontest* function from the *AER* package. The resulting dispersion ratio (1.057) and non-significant p-value (*p*-value = 0.252) confirmed that the variance of the response variable did not significantly exceed the mean, validating the use of the Poisson GLM. This method models the biological relationship between virus richness count data as a linear response with ecological or host factors as the predictor variables affecting virus diversity. Using viral richness as the response variable, multiple model structures were initially explored via Akaike Information Criterion (AIC)^66^, with the final model including mosquito species, collection site, season, and collection year as predictors. To quantify the relative importance of these drivers, we performed an Analysis of Deviance on the selected generalised linear model. The proportion of total deviance explained by each predictor variable was calculated by dividing the deviance reduction attributable to each variable by the Null Deviance. The remaining variation was categorized as unexplained (residual) deviance. Statistical significance of the predictors was determined using Chi-squared tests (χ2), and results were visualized using a pie chart to illustrate the contribution of each predictor variable to the overall variation in viral richness.

### PERMANOVA analysis of community drivers of virome composition

To quantify the drivers of viral community composition, we conducted Permutational Multivariate Analysis of Variance (PERMANOVA) using the *adonis2* function in the *vegan* package. The analysis was performed on a Jaccard distance matrix derived from viral species presence-absence data, which included only viruses detected in at least three individual sequencing libraries. We employed a hierarchical model structure to partition the variance across taxonomic and spatial levels. The model nested mosquito species within their respective genera and collection sites within their respective countries (Country / Site). Additional ecological and temporal factors, including season and collection year, were included as fixed effects. The relative contribution of each predictor variable was determined by the coefficient of determination (R^2^), representing the proportion of total community variation explained.

### Principal coordinates analysis (PCoA) on virome composition

To visualize the influence of host taxonomy and ecological factors on viral community structure, Principal coordinates analysis (PCoA) was performed using the *cmdscale* function. Dissimilarity between samples was calculated using Jaccard distance matrices, which are appropriate for viral presence-absence data. To assess the relative impact of various ecological drivers, we generated a multi-panel ordination series visualizing the interaction between host taxonomy (Genus and Species) and environmental variables (Country and Season). The first two principal coordinate axes were extracted, and the percentage of total eigenvalues explained by each axis was reported. Data were processed and visualized using the *vegan*, *ggplot2*, and *gridExtra* packages, utilising ColorBlind-friendly Viridis scales to ensure clarity and accessibility.

To evaluate multivariate community dispersion, we calculated the distance of each sample to its group centroid using the *betadisper* function based on the Jaccard matrix. Root-mean-square deviation (RMSD) from centroids was computed for each group level across five metadata factors: mosquito genus, mosquito species, country, season, and site. Statistical significance for differences in group dispersion was determined using permutation tests with 999 permutations.

### Indicator species analysis

To identify specific viruses significantly associated with particular host and ecological variables, we performed an Indicator Species Analysis using the *multipatt* function from the *indicspecies* package. This analysis utilized the IndVal.g index (to account for unequal group sizes), with statistical significance determined via 999 permutations. To eliminate low-prevalence noise, mosquito species with at least three samples and viruses detected in at least three sequencing libraries were included in this analysis. Viral indicators were considered significant at *P*-value < 0.05 and visualized in a dot plot to highlight host-specialist and site-specific associations. Mosquito species or categories missing from the final plot either lacked the baseline sample size needed for analysis or carried viruses shared with other local hosts, meaning they lacked a unique viral signature.

### Virus similarity network analysis

Network nodes were constructed by aggregating virus presence-absence data across groups of sequencing libraries sharing combinations of mosquito species, mosquito genus, collection year, season, collection site, and country. Pairwise virome similarity between nodes was quantified using the Jaccard similarity coefficient^67^. The network was visualized as an undirected graph using the *igraph* package, where edges represent the degree of viral species sharing between host populations. To focus on ecologically meaningful relationships while preserving geographical representation, we implemented a hybrid filtering approach: edges were retained if they exceeded a Jaccard similarity threshold of 0.05. The network layout was generated using the Fruchterman-Reingold force-directed algorithm with 5000 iterations to achieve optimal node positioning^68^. Node size was scaled by viral species richness, while edge thickness and transparency were scaled by the strength of Jaccard similarity. Visualizations were rendered using *ggplot2* and *ggrepel*, with nodes labelled by host mosquito and collection site.

### Bipartite network analysis of Madagascar mosquito viromes

To evaluate host-virus associations specifically within Madagascar, we constructed a bipartite network using the *ggraph* package. Nodes were categorized into two distinct types: host mosquito species and viral species. Edges were established based on viral presence, with viral nodes further classified as ’generalists’ (associated with ≥2 host genus) or ’specialists’ (associated with one host genus). The network was visualized using a diagonal bipartite layout, with node size reflecting relative degree centrality.

### Analysis of quantitative abundance (RPM)

To evaluate the distribution and relative abundance of the core virome, the dataset was filtered to isolate the top 50 viral species globally based on cumulative Reads Per Million (RPM). For host-specific and spatial analyses, mean normalized RPM values were calculated for each viral species across mosquito host species and collection sites, respectively. Abundance data were log-transformed using log_10_ (mean RPM + 1). Hierarchical clustering heatmaps incorporating dendrograms for both rows (viral species) and columns (host species or sampling sites) were generated using the *ComplexHeatmap* package in R.

## DATA AVAILABILITY

Raw meta-transcriptomics sequencing reads were deposited in the Sequence Read Archive under BioProject PRJNA1501385. Assembled hallmark gene-coding viral contig sequences have been deposited on GenBank database. The full of identified viral contigs (2290 viral contigs, 966 vOTUs), the filtered dataset for statistical analyses (1401 viral contigs, 263 vOTUs), R code, virus super-clades and mosquito *cox1* phylogenetic trees are deposited in the GitHub repository (https://github.com/anabut123/PREEMPT).

## ACKNOWLEDGEMENTS

This work was funded by the Defense Advanced Research Projects Agency PREEMPT program managed by Dr. Rohit Chitale and Dr. Kerri Dugan [Cooperative Agreement HR001118S0017] to M.-C.S. (the content of the information does not necessarily reflect the position or the policy of the U.S. government, and no official endorsement should be inferred), and by the Laboratoire d’Excellence: Integrative Biology of Emerging Infectious Diseases program (grant ANR-10-LABX-62-IBEID) to C.K. and M.-C.S., by the Agence Nationale de la Recherche (grant ANR-22-CE35-0001, MAMMAMIA) and the Fondation iXcore - iXlife - iXblue Pour La Recherche to M.-C.S.. We thank members of the Viruses and RNA Interference Unit, Josée Dussault, Juan Carlos Ocampo, and Caroline Zanchi for valuable insightful discussions throughout the study. We are grateful to the members of medical entomology teams at Institut Pasteur Bangui, Institut Pasteur Cambodia, Institut Pasteur Madagascar, and Institut Pasteur in French Guiana for undertaking field work and we acknowledge the local communities for their support.

## AUTHOR CONTRIBUTIONS

Conceptualization: C.K., L.F., A.B., M.-C.S.

Methodology: C.K., L.F., H.B., A.B.

Sample collection and processing: C.N., S.B., P.D., N.G., R.G., J.-B.D., C.K., H.B., A.G., A.H.-L.

Data analysis and illustration: A.B., C.K. Writing–original draft: C.K. and A.B.

Writing–review and editing: L.F., A.B., C.N., S.B., P.D., N.G., R.G., J.-B.D., R.C., M.-C.S.

Funding acquisition: C.K. and M.-C.S. Supervision: C.K. and M.-C.S.

## COMPETING INTERESTS

The authors declare no competing interests.

## ADDITIONAL INFORMATION

## Extended data

**Extended Data Figure 1.**
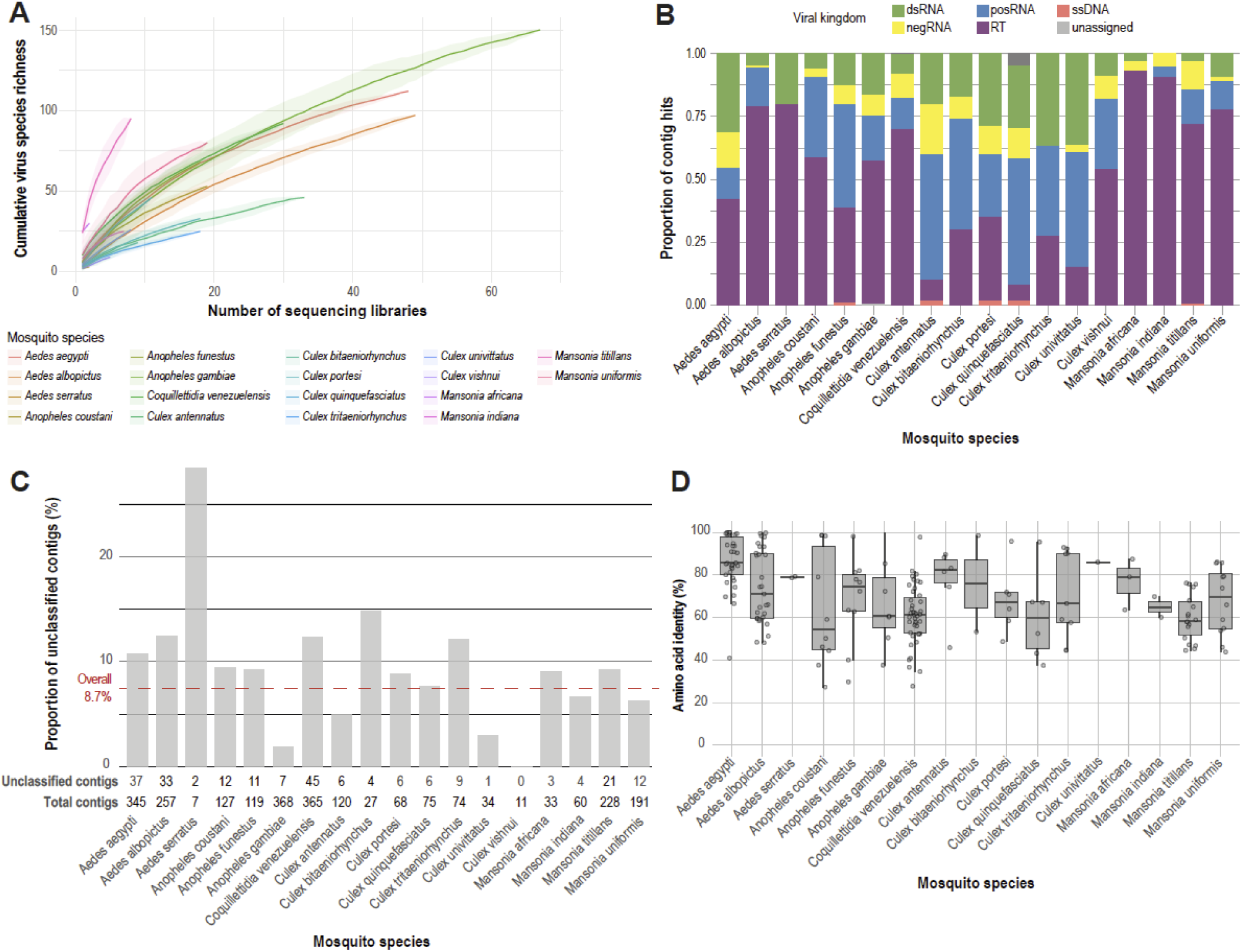
Mosquito virus communities are composed of diverse viral phylogenies, including unclassified viruses. **a**, Rarefaction curves of virus species richness discovered by meta-transcriptomics sequencing of 18 mosquito species. Shading represents 95% confidence intervals. **b,** Relative prevalence of viral kingdoms in each mosquito species. **c,** Proportion of unclassified viruses. Red dashed line denotes overall average of 8.7%. **d,** Amino acid sequence identities of unclassified viral contigs.

**Extended Data Figure 2.**
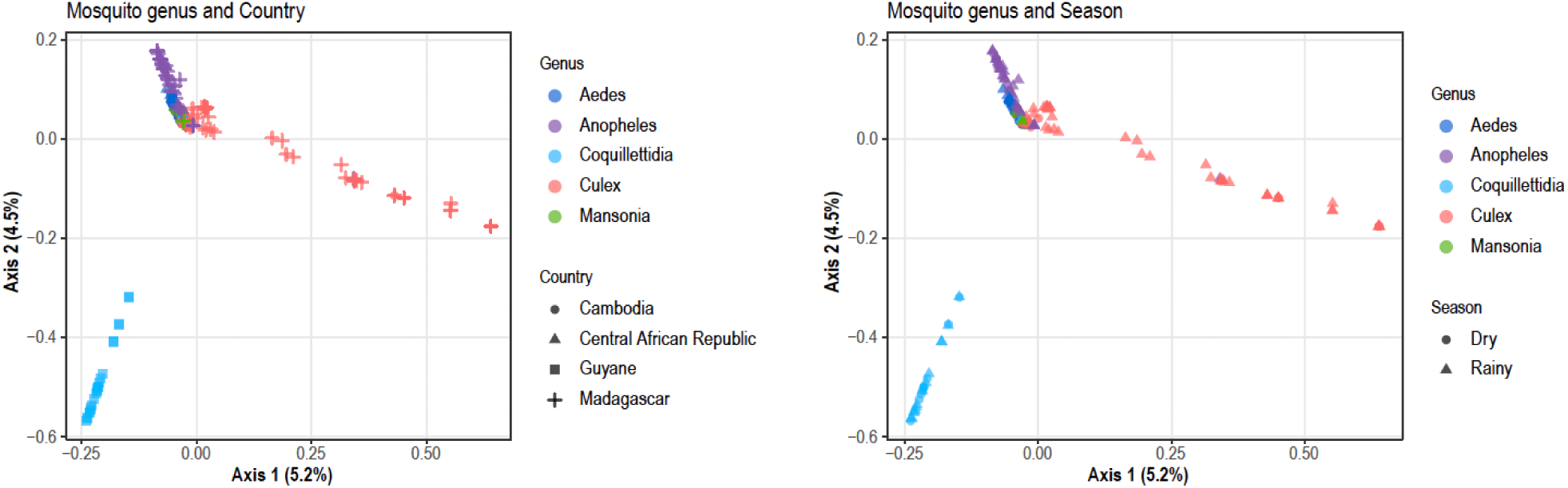
Principal coordinates analysis plots based on a Jaccard distance matrix of virome composition across mosquito populations. Each datapoint represents a mosquito population, where colours denote mosquito genus and shapes denote country or season.

**Extended Data Figure 3.**
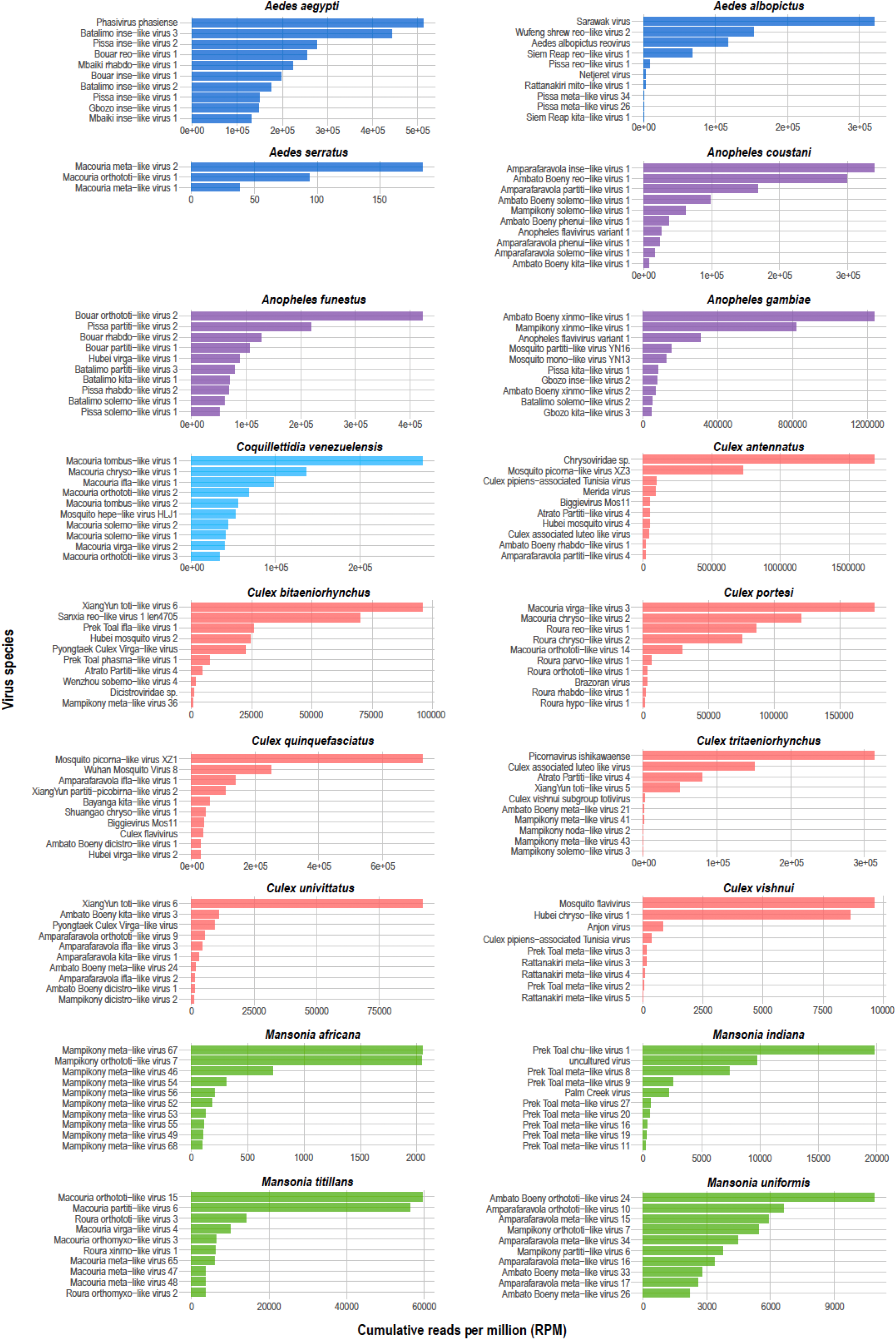
Most abundant viruses per mosquito species. Cumulative reads per million is calculated for each virus across all sequencing libraries per mosquito species and ranked in descending order. Only the ten must abundant viruses are shown.

**Extended Data Figure 4.**
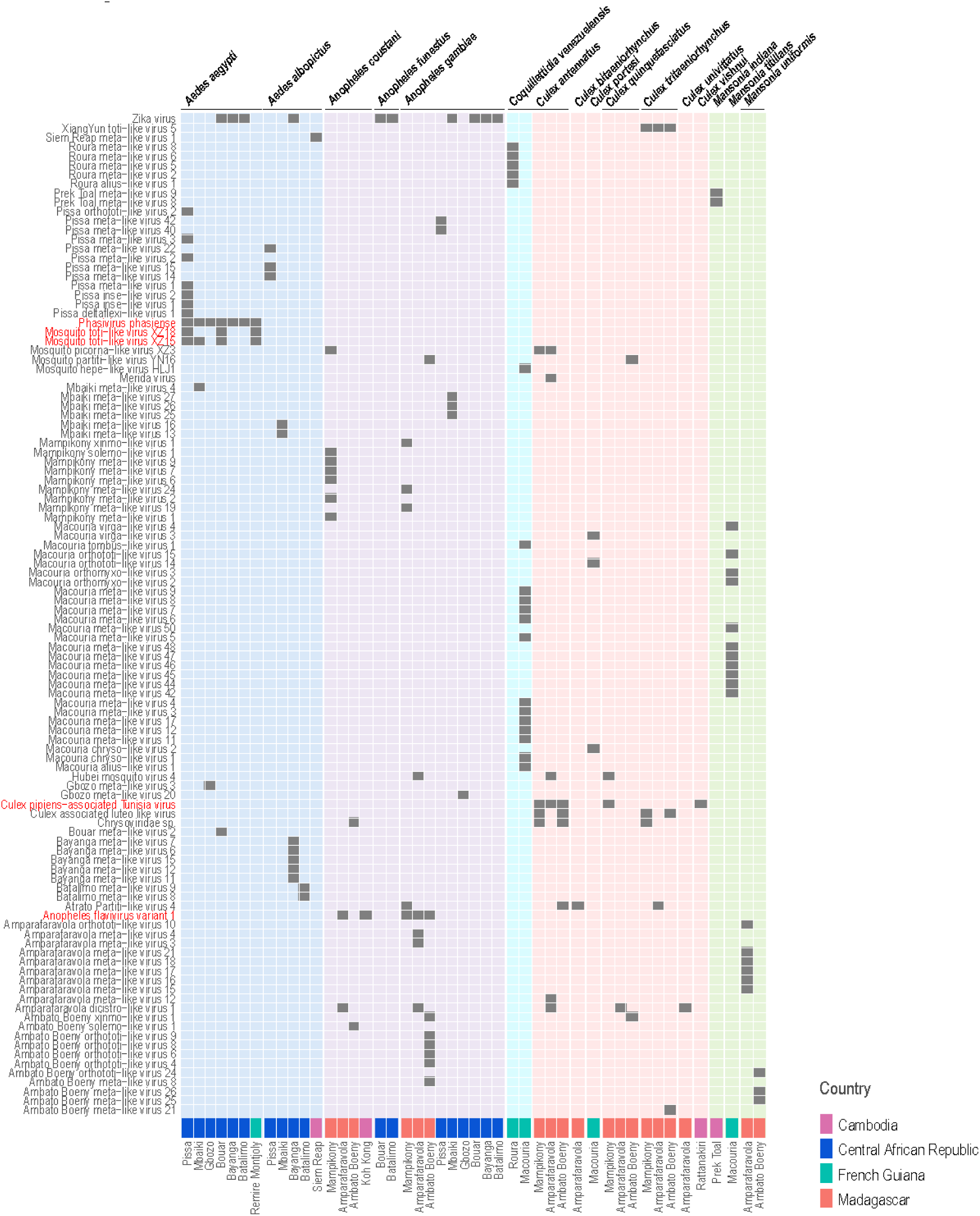
Virus distribution across host and spatial factors. Virus occurrence across mosquito species and geographies (sites and countries). Heatmap background colour corresponds to mosquito genus.

## Supplementary information

**Supplementary Figure 1.**
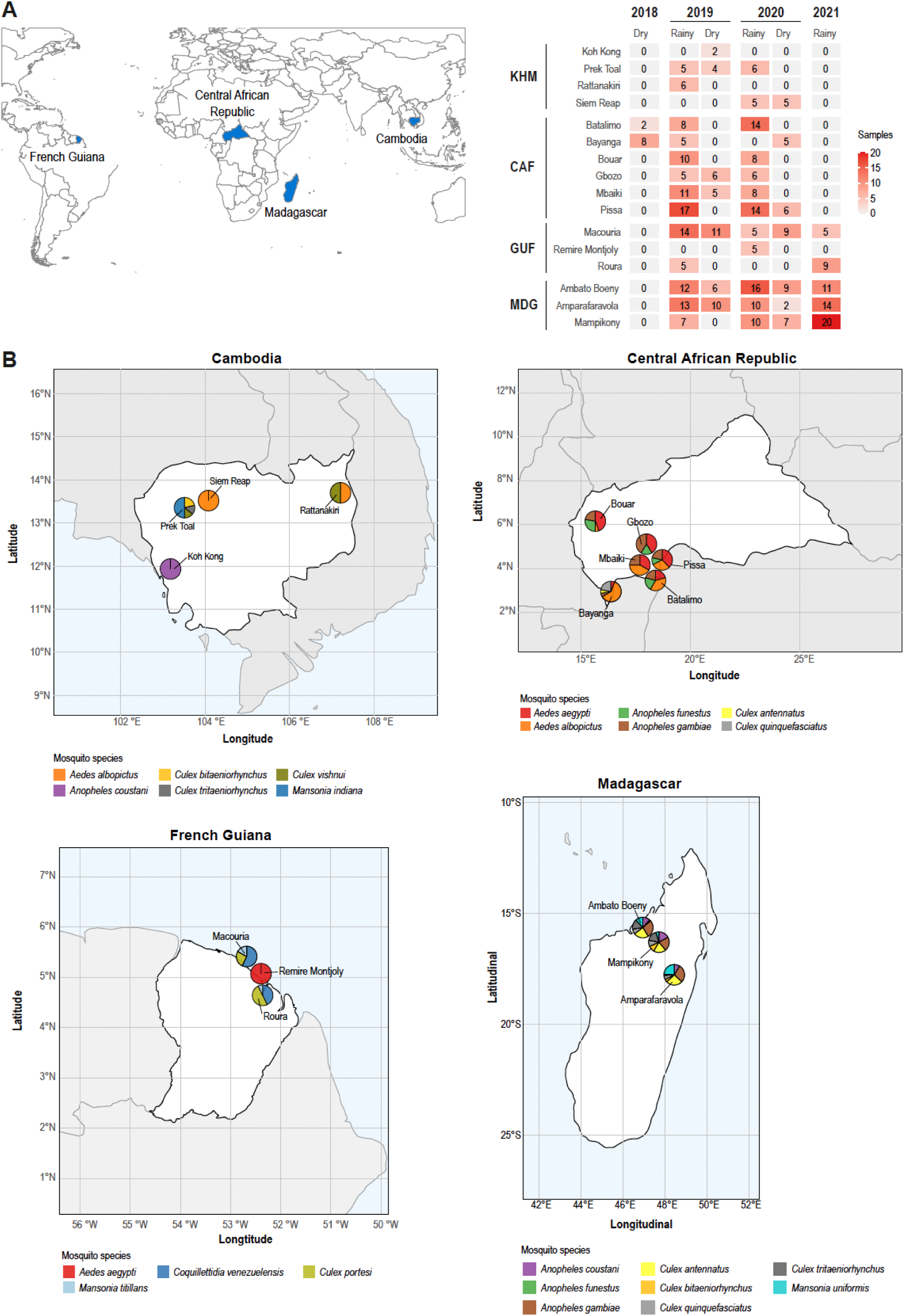
Depth and diversity of mosquito sampling. **a**, Sampling depth per collection site. **b,** Diversity of mosquito species sampled in each site by country.

**Supplementary Figure 2.**
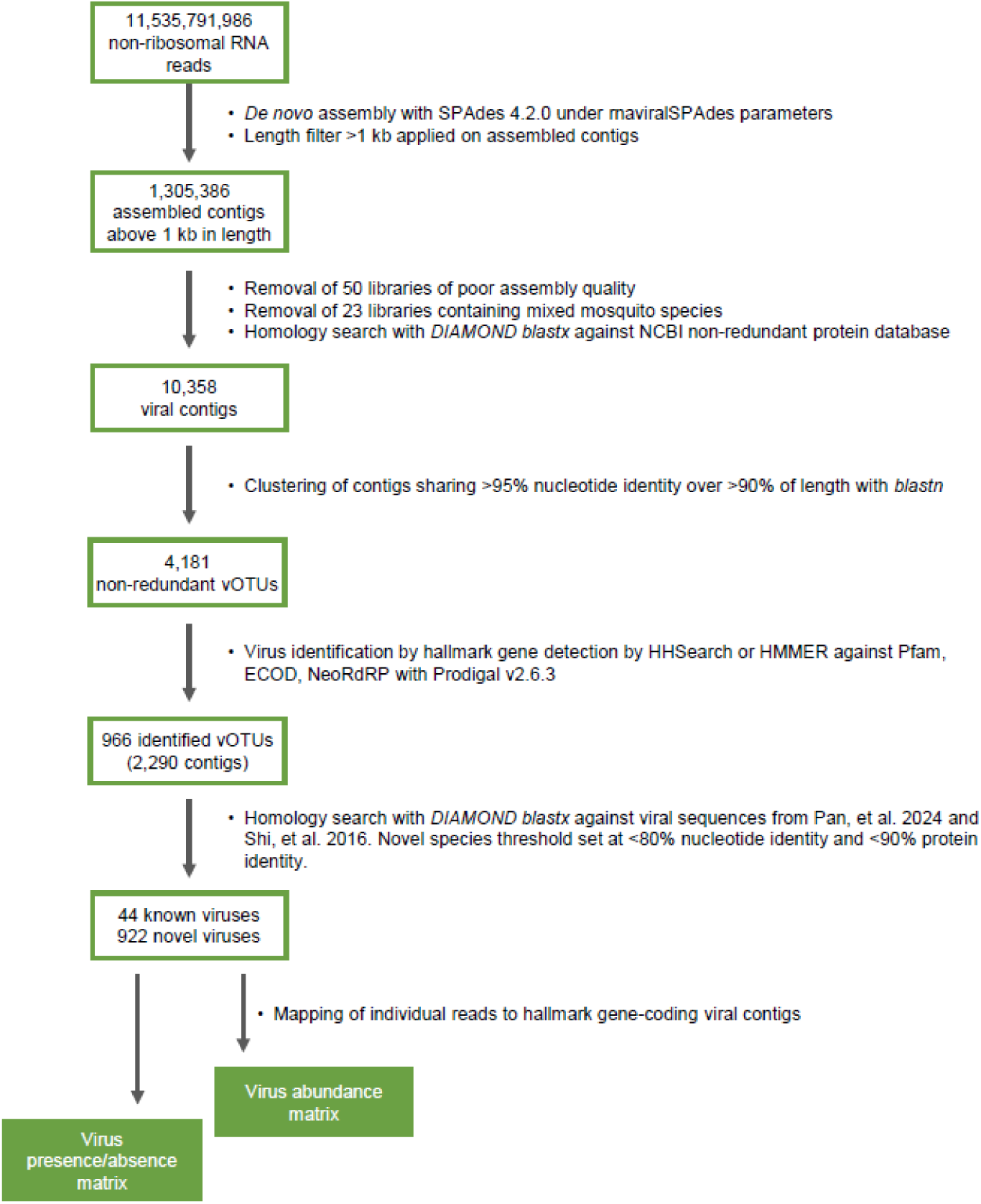
Schematic of bioinformatics workflow for virus detection. Novel viruses were identified through homology search against viral sequences from two landmark studies on invertebrate viromes^2,57^.

**Supplementary Figure 3.**
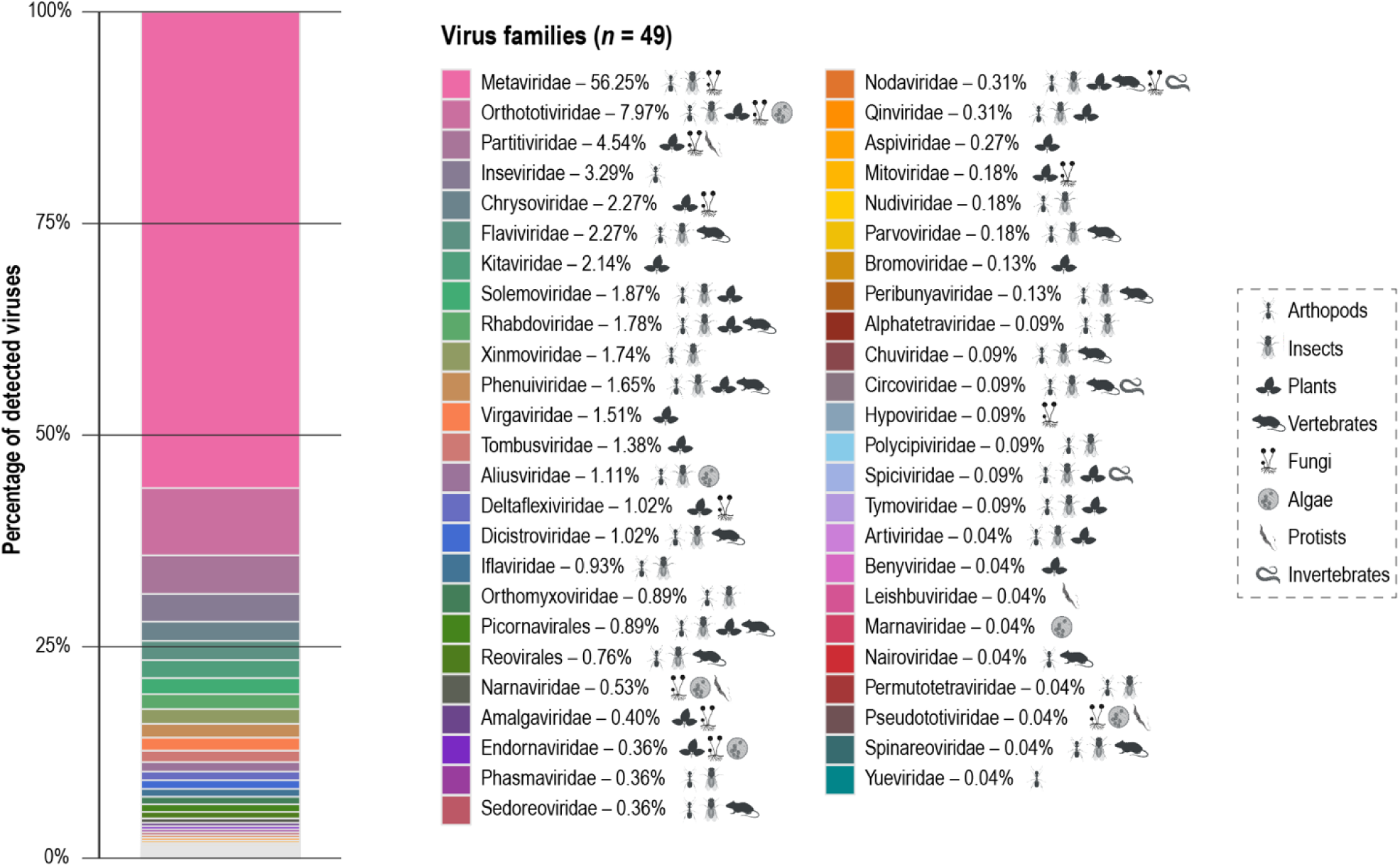
Detected virus families and their host associations. Icons next to each virus family indicate associated hosts based on ICTV information.

**Supplementary Table 1.** Sampling and metadata information of sequencing libraries, each a pool of five female mosquitoes from the same species, site, season and year. (.XLSX file)

**Supplementary Table 2.** List of detected viral contigs. (.XLSX file)

**Supplementary Table 3.**
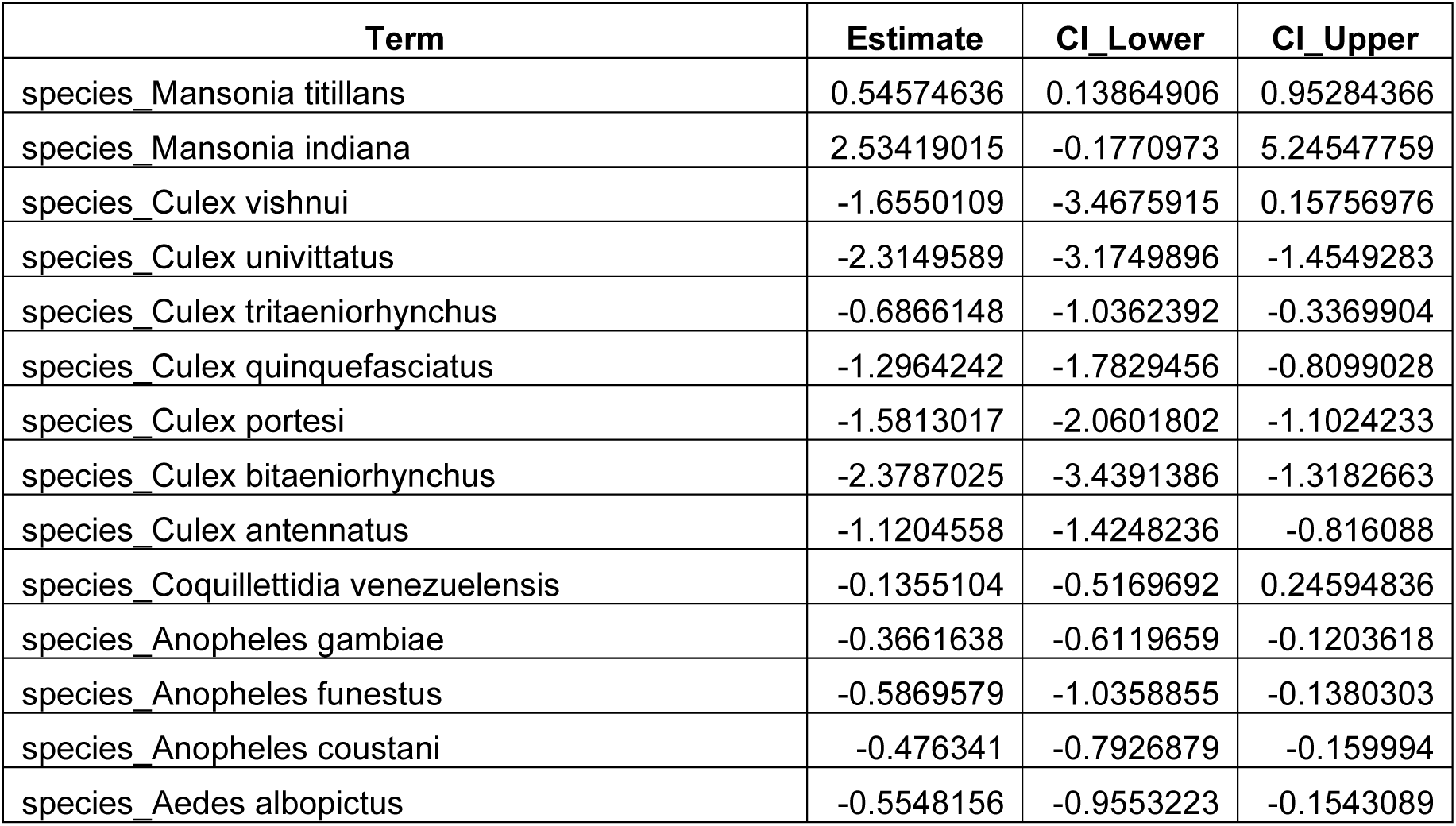
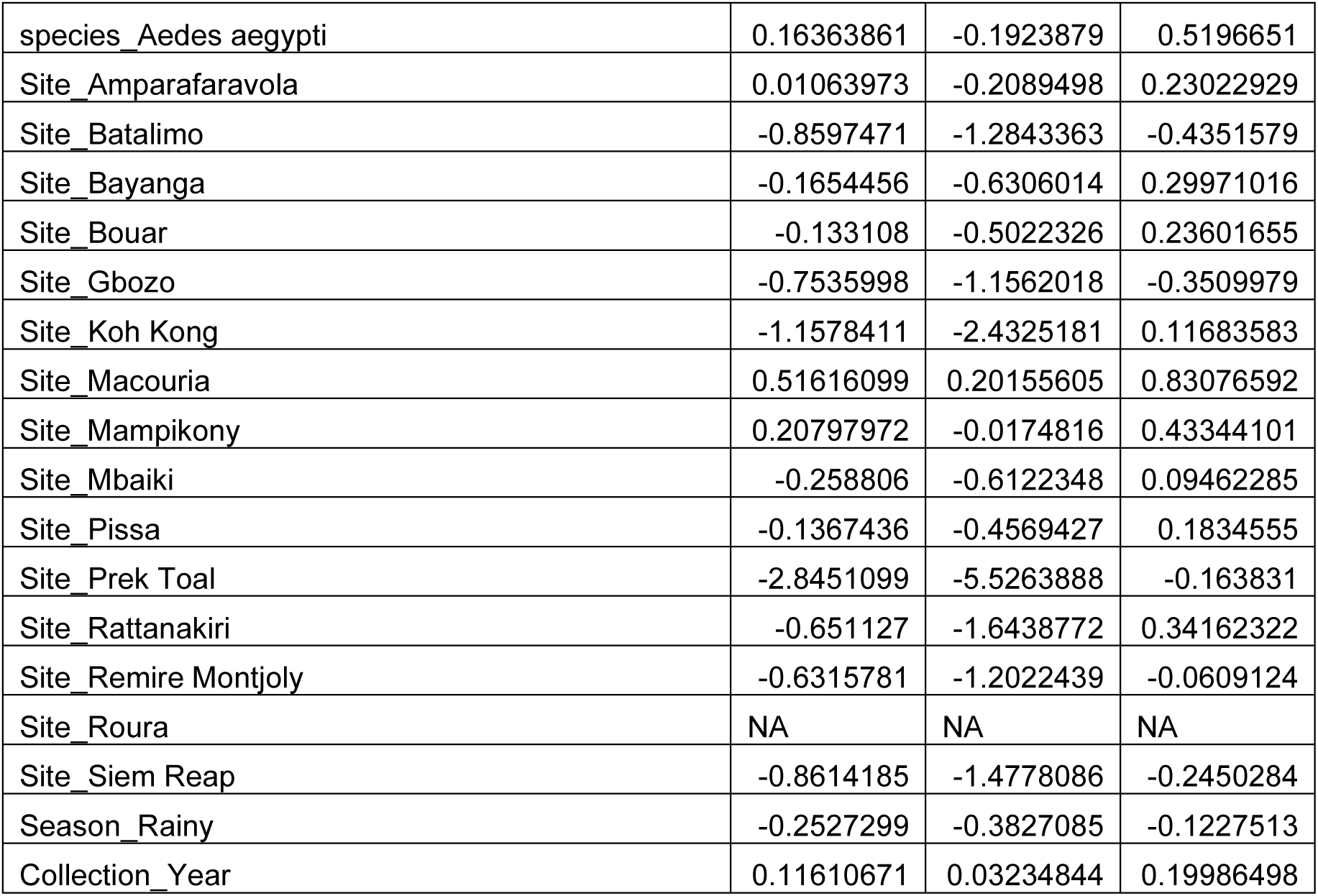
Coefficient estimates and confidence intervals of the best generalised linear model estimating variation in virus diversity.

**Supplementary Table 4.** Root mean squared deviation (RMSD) from centroids by group.

| Variable | Group / Category | RMSD from centroid |
| --- | --- | --- |
| Mosquito genus | Aedes | 0.6928 |
| Mosquito genus | Anopheles | 0.6892 |
| Mosquito genus | Coquillettidia | 0.6562 |
| Mosquito genus | Culex | 0.6807 |
| Mosquito genus | Mansonia | 0.6585 |
| Mosquito species | Aedes aegypti | 0.6727 |
| Mosquito species | Aedes albopictus | 0.6845 |
| Mosquito species | Anopheles coustani | 0.6500 |
| Mosquito species | Anopheles funestus | 0.6710 |
| Mosquito species | Anopheles gambiae | 0.6754 |
| Mosquito species | Coquillettidia venezuelensis | 0.6562 |
| Mosquito species | Culex antennatus | 0.6166 |
| Mosquito species | Culex bitaeniorhynchus | 0.2628 |
| Mosquito species | Culex portesi | 0.6258 |
| Mosquito species | Culex quinquefasciatus | 0.6516 |
| Mosquito species | Culex tritaeniorhynchus | 0.5649 |
| Mosquito species | Culex univittatus | 0.6044 |
| Mosquito species | Culex vishnui | 0.0000 |
| Mosquito species | Mansonia indiana | 0.4811 |
| Mosquito species | Mansonia titillans | 0.5519 |
| Mosquito species | Mansonia uniformis | 0.6035 |
| Country | Cambodia | 0.6686 |
| Country | Central African Republic | 0.6970 |
| Country | Guyane | 0.6842 |
| Country | Madagascar | 0.6903 |
| Season | Dry | 0.6969 |
| Season | Rainy | 0.7017 |
| Site | Ambato Boeny | 0.6734 |
| Site | Amparafaravola | 0.6729 |
| Site | Batalimo | 0.6543 |
| Site | Bayanga | 0.5930 |
| Site | Bouar | 0.6620 |
| Site | Gbozo | 0.6414 |
| Site | Koh Kong | 0.4222 |
| Site | Macouria | 0.6738 |
| Site | Mampikony | 0.6630 |
| Site | Mbaiki | 0.6626 |
| Site | Pissa | 0.6770 |
| Site | Prek Toal | 0.4811 |
| Site | Rattanakiri | 0.5792 |
| Site | Remire Montjoly | 0.5046 |
| Site | Roura | 0.5855 |
| Site | Siem Reap | 0.6447 |

## Notes

### Competing Interest Statement

The authors have declared no competing interest.

### Summary of Updates

New bioinformatics workflow implemented and statistical analysis on drivers of virus diversity and virome composition now performed at virus-species level.

